# Representations of stimulus meaning in the hippocampus

**DOI:** 10.1101/2024.10.14.618280

**Authors:** Jeremy S. Biane, Max A. Ladow, Austin Fan, Hye Sun Choi, Lexi Zichen Zhou, Shazreh Hassan, Daniel L. Apodaca-Montano, Andrew O. Kwon, Joshua X. Bratsch-Prince, Mazen A. Kheirbek

## Abstract

The ability to discriminate and categorize the meaning of environmental stimuli and respond accordingly is essential for survival. The ventral hippocampus (vHPC) controls emotional and motivated behaviors in response to environmental cues and is hypothesized to do so in part by deciphering the positive or negative quality of these cues. Yet, what features of the environment are represented in the activity patterns of vCA1 neurons, and whether the positive or negative meaning of a stimulus is present at this stage, remains unclear. Here, using 2-photon calcium imaging across six different experimental paradigms, we consistently found that vCA1 ensembles encode the identity, sensory features, and intensity of learned and innately salient stimuli, but not their overall valence. These results offer a reappraisal of vCA1 function, wherein information corresponding to individual stimulus features and their behavioral saliency predominates, while valence-related information is attached elsewhere.

## INTRODUCTION

Brains have evolved to take sensory information from the environment and generate appropriate behavioral and physiological responses. Certain sensory stimuli have innate meaning to an organism, whereas other stimuli become robust triggers of emotional responses and/or behavioral outputs only through associative learning (Biane et al., 2023; Fanselow and Dong, 2010; Jimenez et al., 2018; Kheirbek et al., 2013; Komorowski et al., 2009; Royer et al., 2010; Strange et al., 2014; Turner et al., 2022). In each case, the brain must recognize the stimulus in question – i.e. encode its identity – and simultaneously attach importance to it.

As the neuroanatomy and circuit basis of emotional processing is mapped with increasing resolution, the ventral hippocampus (vHPC) stands out as a central hub and an interface between emotional and cognitive processing (Turner et al., 2022). Consistently over the past 30 years, numerous studies have shown that lesioning, inhibiting, or stimulating the vHPC can modulate defensive behaviors or reward-related approach behaviors (Bannerman et al., 2003; Britt et al., 2012; Ciocchi et al., 2015; Henke, 1990; Jacobson and Sapolsky, 1991; Jimenez et al., 2018; Kjelstrup et al., 2002; Komorowski et al., 2013; LeGates et al., 2018; Padilla-Coreano et al., 2019, 2016; Parfitt et al., 2017; Reed et al., 2018; Royer et al., 2010). These observations suggest that the vHPC is crucial for capturing and representing aversive and rewarding features of an environment – that is, the valence of those particular features – and orchestrating physiological and behavioral responses. Yet, fundamental properties of this region remain unresolved. Perhaps chief among these is how salient stimuli are represented and processed by the vHPC as it contributes to selecting appropriate emotional and behavioral responses.

Recent studies using optogenetic manipulations, in vivo recordings, and immediate early gene immunohistochemistry suggest that discrete classes of ventral CA1 (vCA1) pyramidal neurons represent either rewarding or aversive information, and control distinct aspects of behavior (Britt et al., 2012; Ciocchi et al., 2015; Jimenez et al., 2020, 2018; LeGates et al., 2018; Okuyama et al., 2016; Padilla-Coreano et al., 2019; Shpokayte et al., 2022; Turner et al., 2022; Wee et al., 2024; Xu et al., 2016; Yang et al., 2020). As many vCA1 projection neurons send axons to a single downstream brain region (Gergues et al., 2020), it has been further inferred that distinct parallel pathways from vCA1 carry discrete types of information. Based on these studies, recent hypotheses posit that vCA1 incorporates information regarding the valence of events in a labeled-line-like organization, with overlapping ensembles of neurons encoding stimuli of similar valence. In this scenario, a key role of vCA1 is to encode the positive or negative valence of stimuli.

On the other hand, vCA1 representations of salient environmental cues has been shown to be relatively stable(Yun et al., 2023), even when the outcomes associated with such cues is altered(Biane et al., 2023), calling into question prevailing interpretations of vHPC function. Deciphering what information is extracted by vCA1 ensembles would provide a central element of vHPC function, and by extension, emotional processing. Thus, we set out to unravel the logic of emotional processing in the vHPC by directly testing whether the positive or negative meaning of a stimulus is represented and, more broadly, elucidating what information is embedded within vCA1 activity.

## RESULTS

### Coding of innately salient stimuli in vCA1

We first set out to understand the logic by which vCA1 represents stimuli that are innately aversive or rewarding. Previous work has shown that ensembles of vCA1 neurons respond to a wide range of unconditioned stimuli of different modalities and of both positive and negative valence, including sucrose rewards, footshocks, anxiogenic environments, social stimuli, and meaningful olfactory stimuli (Adhikari et al., 2010; Biane et al., 2023; Ciocchi et al., 2015; Jimenez et al., 2020, 2018; Okuyama et al., 2016; Padilla-Coreano et al., 2016; Reed et al., 2018; Shpokayte et al., 2022; Taxidis et al., 2020; Turner et al., 2022). Our first question was whether positive and negative stimuli induce separable response patterns consistent with the generation of a valence code in vCA1 (Gergues et al., 2020; Tannenholz et al., 2014; Xia and Kheirbek, 2020). To ask this, we injected adeno-associated virus (AAV) expressing the calcium-sensitive protein GCaMP6f (Chen et al., 2013) into vCA1 and implanted a gradient index lens (GRIN) above this region. We then used 2-photon microscopy to image fields of view (FOV) of vCA1 as mice were exposed to a panel of salient stimuli of differing identities and valences (Fig. 1**A-C**).

**Figure 1.**
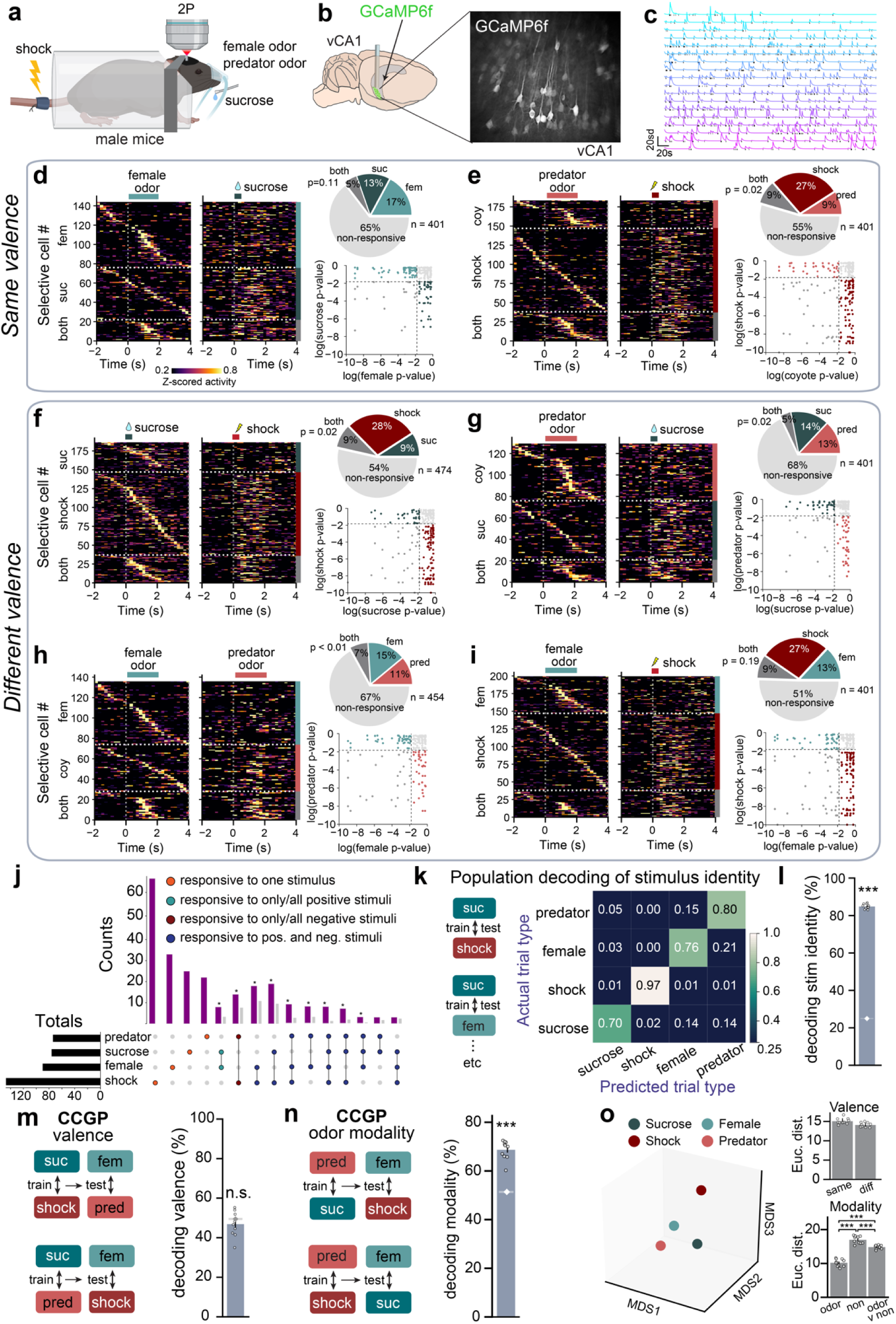
vCA1 responses to appetitive and aversive stimuli. **a,** Study design. Mice were exposed to 2 odors (predator and female), sucrose rewards and shocks (See Methods for details). **b**, Left: schematic of virus injection and GRIN lens implant site. Right: field of view of vCA1 neurons expressing GCaMP6f imaged through a 2-photon microscope. **c**, Example traces from a population of vCA1 neurons. Ticks below the traces indicate event times. **d-i,** Each panel shows the single-cell responses in a pairwise analysis of each stimulus pair. Heatmaps plot all cells significantly responsive to either of the two stimuli (stim 1 trials plotted on left, stim 2 trials on right). Stim 1 heatmaps are sorted with respect to time of greatest activity, and stim 2 heatmaps are sorted with respect to stim 1. horizontal dashed lines demarcate cells responsive to stimulus 1 (top), stimulus 2 (middle), or both stimuli (bottom). Pie charts take into account all cells, computing the percentage that are responsive to each stimulus, both stimuli or none. P-values reflect the observed proportion of cells responsive to both stimuli vs that expected by chance (see Methods). Scatter plots: we computed each cell’s stimulus responsivity P-value (see Methods). Cells inside both dashed lines exceeded the significance threshold (p = 0.05) for both stimuli. **j,** Upset plot for all stimulus-responsive neurons (n = 249). Purple bars indicate the true number of cells and grey bars the number expected by chance. **k**, Left: schematic of decoding paradigm. Right: confusion matrix for decoding the 4 stimuli. Accurate decoding is along the diagonal. **l,** Decoding stimulus identities was significantly greater than chance in vCA1. Individual dots represent the results of a single decoding run. White diamond and horizontal line represent mean +/− s.e.m. chance decoding accuracy. **m,** Decoding valence in vCA1. The decoder was trained on one pair of appetitive vs aversive stimuli and tested on the complementary pair. Decoding accuracy was no better than chance, suggesting that there was no grouping of these stimuli by valence in vCA1. **n,** Decoding stimulus modality (odor vs non) in vCA1. The decoder was trained on one pair of odor vs non-odor stimuli, and tested on the complementary pair. Stimulus modality could be decoded significantly better than chance. **o,** MDS visualization, where the relationship between activity patterns is represented in geometrical space; the closer two points are in space, the more similar their activity patterns are. We could not detect a clustering of stimulus representations with similar valences (the mean of 10 MDS runs plotted to the right). In contrast, odor representations were significantly closer to one another compared to the distance between sucrose and shock, or the distance between odor and non-odor representations. Pairwise breakdown of Euclidean distances between each valence group is presented in Supplementary Fig 2h. \*\*\**P* < 0.001; \*\**P* < 0.01, \**P* < 0.05. All error bars indicate mean ± s.e.m. See Supplementary Table 1 for all statistical analysis details.

We imaged population activity in vCA1 in a four-stimulus design, where two positive stimuli (sucrose rewards and female odor) and two negative stimuli (mild tail shocks and predator (coyote) odor) were presented individually to thirsty male mice (Fig. 1**A**). Each stimulus robustly activated vCA1, evoking activity in roughly 20-35% of recorded neurons. While many stimulus-responsive neurons responded to only a single assayed stimulus, ∼40% of these neurons responded to two or more stimuli (Fig. 1**J**).

The key question for us was whether stimuli of the same valence (e.g., sucrose and female urine) recruited overlapping ensembles of neurons, while stimuli of opposite valence activated separate ensembles. Strikingly, however, the propensity of neurons to respond to stimuli of the same valence was no greater than their likelihood of responding to stimuli of opposite valences. Indeed, looking solely at multi-responsive cells, only a small percentage responded selectively to positive (8%) or negative stimuli (14%); the vast majority (78%) being responsive to stimuli across valences. This suggests that responsive neurons mainly responded to just one stimulus, but critically neurons responsive to multiple stimuli do so in a way that is largely independent of the valences of the stimuli.

We corroborated these results using microendoscopy recordings from vCA1 in freely moving mice (Supplementary Fig. 1). Previous work identified a subpopulation of vCA1 neurons preferentially active in the anxiogenic open arms of the elevated plus maze (EPM) (Ciocchi et al., 2015; Jimenez et al., 2018). Here, we asked whether these “anxiety cells” were also more likely to respond to additional negative stimuli of different modalities.

For this, we imaged mice in the EPM, then imaged the same field of view when mice were presented with a panel of positive, negative, and neutral odors, as well as sucrose and footshocks. We did not find evidence that open arm cells were more likely to respond to aversive stimuli than closed arm cells, and *vice versa* (Supplementary Fig. 1). These data suggest that vCA1 neurons active in anxiogenic open environments do not constitute a group of aversion-coding neurons, as they showed no significant preference for responding to other types of aversive stimuli.

Having found no prevalent encoding of stimulus valence at the level of single-cell responses in vCA1, we next asked whether stimulus valence information might be embedded within the overall population response patterns. Consistent with the above single-cell analyses, the identity of each stimulus could be read out from population activity with high accuracy using linear classifiers (Fig. 1**K,L**). To assess valence decoding, we first generated a decoding confusion matrix (Fig. 1**K**), in which the more similar two stimulus representations are, the more likely they are to be confused with one another by the decoder. We examined if responses to stimuli of one valence class (e.g., sucrose and female odor) were more likely to be confused with each other than with responses to stimuli of the opposite class (e.g., footshock and predator odor). This was not, however, the case. Stimulus representations were equally likely to be confused with representations of stimuli belonging to the same or the opposite valence class.

To test whether the appetitive or aversive nature of a stimulus was represented in the population activity in a different way, we assessed the cross-condition generalization performance (CCGP) (Bernardi et al., 2020) for valence by training a linear classifier to discriminate activity between an appetitive trial type (e.g., sucrose) versus an aversive trial type (e.g., shock), then testing this classifier’s “valence decoding” accuracy when using data from the complementary trial types (e.g., female and coyote; see Fig. 1**M**). Here, high decoding accuracy would indicate similar neural representations within trial types of similar valence (e.g., sucrose and female). However, consistent with the above results, CCGP valence decoding performed no better than chance. Exploration of decoding performance using a variety of nonlinear kernels and regularization parameters also failed to find evidence of a valence signal (Supplementary Fig 2). This, again, indicates that neural representations of stimuli of the same valence were no more like one another than they were to those of opposite valence.

Although the confusion plot in Fig. 1**K** showed no convincing evidence of valence encoding, it did show a tendency to miscategorize the two odor stimuli. We, therefore, tested whether a generalized representation of odor was present in the neural activity by training a decoder to discriminate one odor stimulus from one non-odor stimulus, then testing it on the other odor/non-odor stimuli pair (“odor modality decoding”). Here, we found CCGP accuracy for odor modality to be significantly greater than chance (Fig. 1**N**).

These CCGP decoding results for valence and odor modality were corroborated using multidimensional scaling (MDS) to visualize the geometric architecture of the hippocampal representations. We found that the distance between representations of stimuli with similar valences were no closer than those of stimuli with opposite valences (Fig. 1**O**). It also showed that the distance between the two odor representations was significantly shorter than the distance between representations of non-odor stimuli, and between odor and non-odor representations.

Together, these results demonstrate that vCA1 robustly discriminates the identity and modality of sensory stimuli but suggest this brain region does not encode the valence of behaviorally important stimuli (Fig. 1**O**).

### Representations of emotionally salient stimuli are stable across days

In the dorsal hippocampus, the tuning of place cells remains relatively stable upon subsequent exposures to the same context. We thus asked whether such stability was also present in vCA1 for representations of unconditioned stimuli by measuring neuronal activity when the same task was repeated 24 hours later (Fig. 2**A,B**). First, as with our day 1 results, both single-cell and population response properties on day 2 demonstrated that the identity, but not the valence, of stimuli was represented in vCA1 (Figs. 2**C-I** and Supplementary Fig. 3). To assess the stability of stimulus responses across days, we asked whether stimulus-responsive cells from day 1 were activated by the same stimulus on day 2. The proportion of cells responsive to each of the four stimuli across both days was significantly higher than chance (Fig. 2**C-F**). We also observed response stability at the population level, as a decoder trained to discriminate stimulus representations on day 1 performed significantly better than chance when classifying day 2 stimulus representations (Fig. 2**J,K**). Thus, in vCA1, neural representations of unconditioned stimuli display stability across days.

**Figure 2.**
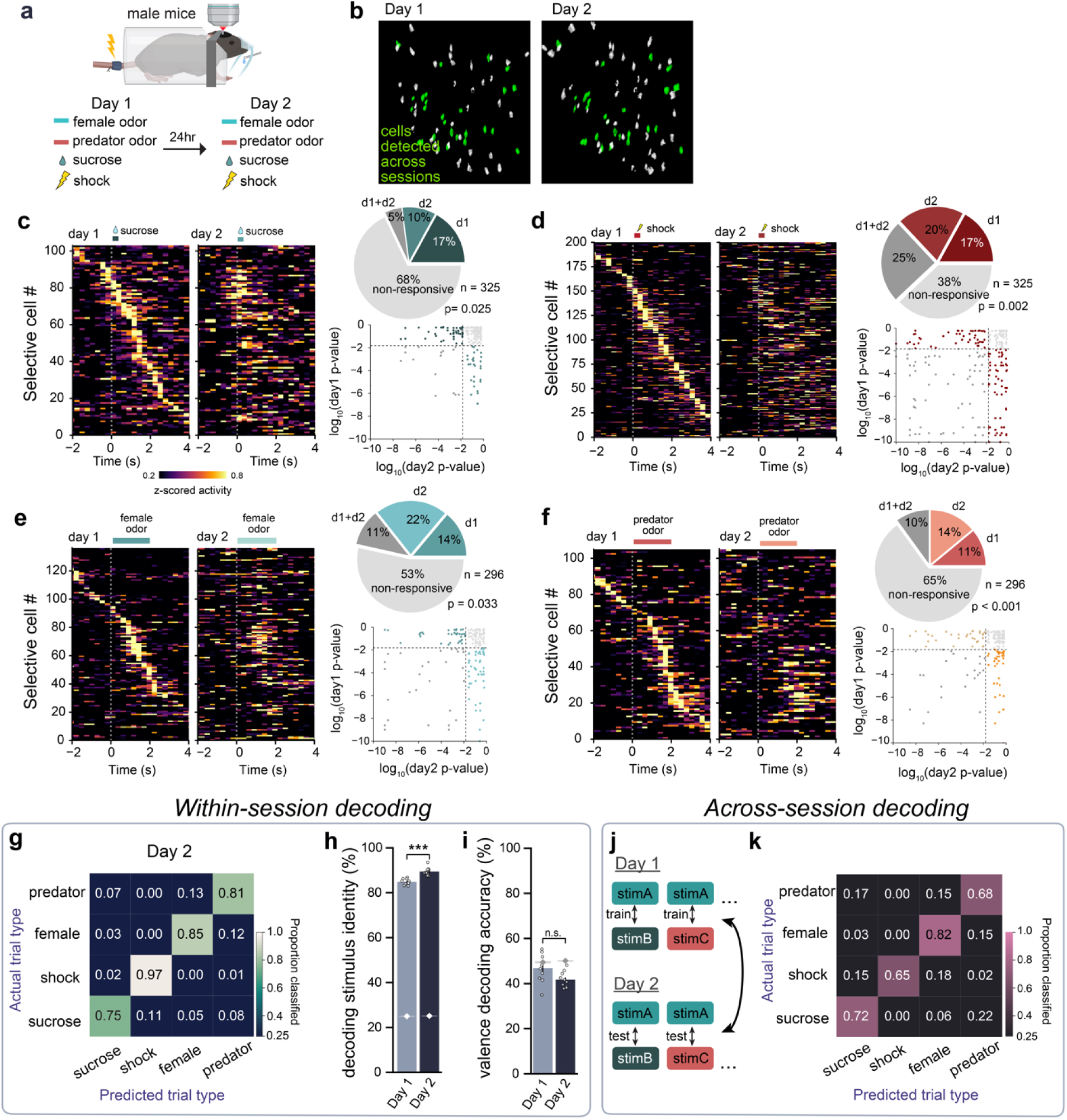
Across day stability of representations of salient stimuli. **a-b,** Experimental design. The same vCA1 field of view was imaged and cells were tracked across two consecutive days to determine the stability of stimulus-responsive vCA1 neurons. **c-f,** We generated heatmaps of normalized activity for cells responsive in Day 1, and examined responses in D2. Pie charts indicate responsive neurons for each stimulus on the 2 days and those that responded in both days, along with p-value testing the significance of the observed proportion of cells active on both days vs the proportion expected by chance. **g,** Confusion matrix for decoding the 4 stimuli on day 2. **h,** Stimulus identity decoding accuracy is better than chance on both days of testing, and increases modestly on D2 of testing. White diamonds and shading indicate mean +/− s.e.m. chance decoding accuracy. **i,** Valence decoding was similar to chance on both days, suggesting that repeated exposure across days did not increase the likelihood that valence would be computed in vCA1. **j,** Cross session decoder to examine stability of stimulus representations. A linear decoder was trained to discriminate each stimulus pair from population activity during one session, and classification accuracy was tested using activity patterns from the other day (performed in both directions, decoding results are average of both). **k,** Confusion matrix for cross session decoding accuracy shows robust cross day stability for the representations of stimulus identity. \*\*\**P* < 0.001; \*\**P* < 0.01, \**P* < 0.05. All error bars indicate mean ± s.e.m. See Supplementary Table 1 for all statistical analysis details.

Finally, in a previous study (Biane et al., 2023) we found the representations of innately neutral odor cues become more distinguishable after animals learn to associate an odor cue with an upcoming reward. We therefore asked if, in addition to being relative stability, the representations of these four innately salient stimuli nevertheless became more distinct with repeated exposures. Here, we also found an increase in decoding accuracy for the identity of unconditioned stimuli across days. This suggests that repeated encounters with salient stimuli enhance stimulus discrimination (Fig. 2**H**), even in the absence of associative learning or explicit training.

### Identity, but not valence, dominates vCA1 representations for stimuli within the same sensory modality

The above results indicate that vCA1 encodes the identity of a stimulus – including its sensory modality – but not its valence. However, as with previous work (Ciocchi et al., 2015; Shpokayte et al., 2022), we have so far compared a battery of stimuli that differed in sensory modality. We thus next used a panel of four liquid gustatory stimuli so that the sensory modality was kept constant. Crucially, in activating four distinct canonical taste pathways, two of these stimuli, sucrose and umami, are appetitive stimuli while the other two, bitter and sour, are aversive (Sinclair et al., 2015; Tordoff et al., 2015) (Fig. 3**A**).

**Figure 3.**
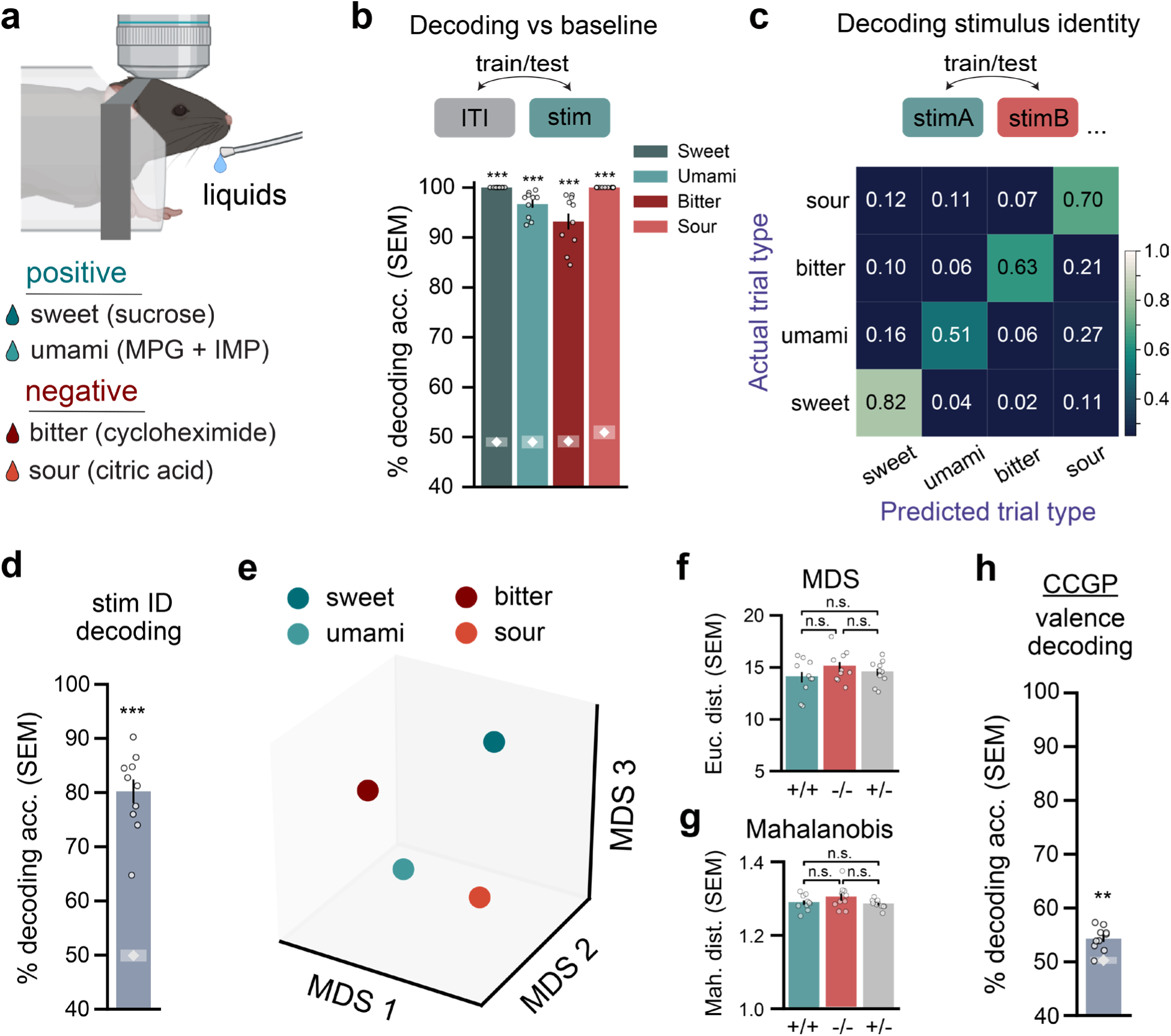
Robust vCA1 encoding of stimulus identity, but not valence, across stimuli of uniform sensory modality. **a,** Study design. vCA1 activity was imaged while mice were exposed to 4 liquids possessing positive or negative innate valence. **b,** Neural activity during liquids consumption could be decoded from baseline (ITI) activity with high accuracy, attesting to the saliency of each stimulus. White diamonds and shading indicate chance decoding mean accuracy ± s.e.m. **c,** Confusion matrix when decoding trial type. Stimulus identity could be accurately decoded. Information about stimulus valence, however, was not evident, as stimuli were typically just as likely to be confused with a stimulus with opposite valence as for a stimulus belonging to the same valence class. **d,** Overall, stimulus identity could be decoded with high accuracy, and well above chance levels. **e,** MDS visualization of similarities among stimulus activity patterns showed a separation of all stimulus patterns, and no clustering of stimuli with shared valence. **f** There was no difference in Euclidean distance between MDS representations for stimuli belonging to the same (+/+, −/−) or different (+/−) valence classes. **g,** Similarly, the Mahalanobis distance did not differ when comparing stimulus representations belonging to the same (+/+, −/−) or different (+/−) valence classes. **h,** CCGP valence decoding (see Fig 1m for schematic) displayed low accuracy but was significantly above chance levels (chance mean = white diamond; chance SEM = line shading). \*\*\**P* < 0.001; \*\**P* < 0.01, \**P* < 0.05. All error bars indicate mean ± s.e.m. See Supplementary Table 1 for all statistical analysis details.

First, we found that the neural activity evoked by each flavored solution could be decoded from background activity with high accuracy, verifying the saliency of each stimulus (Fig. 3**B**). And, as with the cross-modal stimuli, we again found that the identity of each solution could be decoded from vCA1 activity, further supporting the idea that vCA1 encodes the identity of salient, unconditioned stimuli (Fig. 3**C,D**).

We next asked if holding modality constant would allow some kind of valence signal to be detected. However, the confusion matrix shown in Fig. 3**C** displayed no clear pattern of valence encoding. Indeed, three of the four stimuli showed the highest level of confusion with stimuli belonging to the *opposite* valence class. To probe valence encoding further, we again examined the distance between neural representations following MDS (Fig. 3**E,F**), as well as the Mahalanobis distance performed on the full, unreduced dataset (Fig. 3**G**). Neither analysis showed a significant difference in the distance between stimuli belonging to either the same or differing valence class. Finally, CCGP valence decoding accuracy (54.3%) was, in contrast, slightly above chance albeit well below identity decoding accuracy (80.2%) (Fig. 3**J**; Supplementary Fig. 4 shows decoding results when using different nonlinear kernels and regularization parameters, some of which were no better than chance). While this above chance result may reflect a presence of valence encoding within the taste domain, we note that differences in consummatory behavior across valence classes, such as body/orofacial movements, may also contribute to minor differences in hippocampal activity. Overall, however, these findings are consistent with the cross-modal results, namely that vCA1 robustly encodes stimulus identity, but not stimulus valence.

Above, we used four liquid stimuli that each activated a unique taste pathway. To test whether taste pathway itself is represented by vCA1, we presented mice with liquid unconditioned stimuli that recruited overlapping or distinct taste pathways: two stimuli, chocolate and vanilla milk, which both activate sweet taste pathways and two stimuli, quinine and high NaCl, which both activate bitter sensory neurons (Oka et al., 2013). Additionally, in the same session we exposed mice to a positive odor, peanut oil, and a negative odor, isopentylamine (Root et al., 2014), to compare stimuli of differing modalities (Supplementary Fig. 5A). The results mirrored those above. Decoding accuracy was, again, high for stimulus identity. And stimulus modality (here gustatory versus olfactory) could be decoded with high precision. But once more, decoding was indistinguishable from chance for valence (Supplementary Fig. 5C-G).

Interestingly, within the liquid domain, representations of solutions with similar taste qualities tended to be miscategorized with one another when decoded (Supplementary Fig. 5C). This was confirmed using MDS to visualize the geometric architecture of these representations, where we observed that population responses to liquids of the same taste quality (i.e. sweet or bitter) tended to cluster together (Supplementary Fig. 5E).

CCGP for taste quality (Supplementary Fig. 5F,I) further indicated that liquids possessing the same taste quality (e.g., vanilla and chocolate milk) contained overlapping representations which were clearly discriminable from representations of stimuli in the other taste category (high NaCl and quinine; Supplementary Fig. 5E,G). These data suggest that vCA1 activity encodes both the unique identity of gustatory stimuli and the general class to which they belong.

### Stable identity coding following switch of stimulus valence

Given the centrality of the hippocampus to learning and memory, we next asked if a learned reversal of stimulus valence would impact its neural representation in vCA1, or whether the identity of the stimulus would continue to be stably represented, as suggested thus far. To test this, we converted an intrinsically positive liquid treat into a conditioned aversive stimulus. Specifically, mice underwent conditioned taste aversion (CTA), wherein innately appetitive vanilla milk was paired with a lithium chloride injection to induce malaise, resulting in future avoidance of the paired liquid.

In step 1, prior to CTA, neural activity in vCA1 was imaged during delivery of three solutions that differed in their innate valence (Dolensek et al., 2020; Stalnaker et al., 2014; Tordoff et al., 2015): vanilla milk (appetitive), high NaCl (aversive), and low NaCl (appetitive; Fig. 4**A**). This configuration allowed us to assess whether reversing the milk’s valence altered the similarity of its representation relative to those of the two unconditioned positive and negative stimuli. In step 2, CTA was induced and shown to be effective behaviorally (Fig. 4**B**), as mice initially showed a strong preference for milk over water, but virtually avoided milk consumption following CTA. In step 3, one day after CTA administration, vCA1 was again imaged while mice consumed the same three solutions as before. We tracked neurons across the two recording sessions and examined how both population level and single-cell response properties differed pre- and post-CTA.

**Figure 4.**
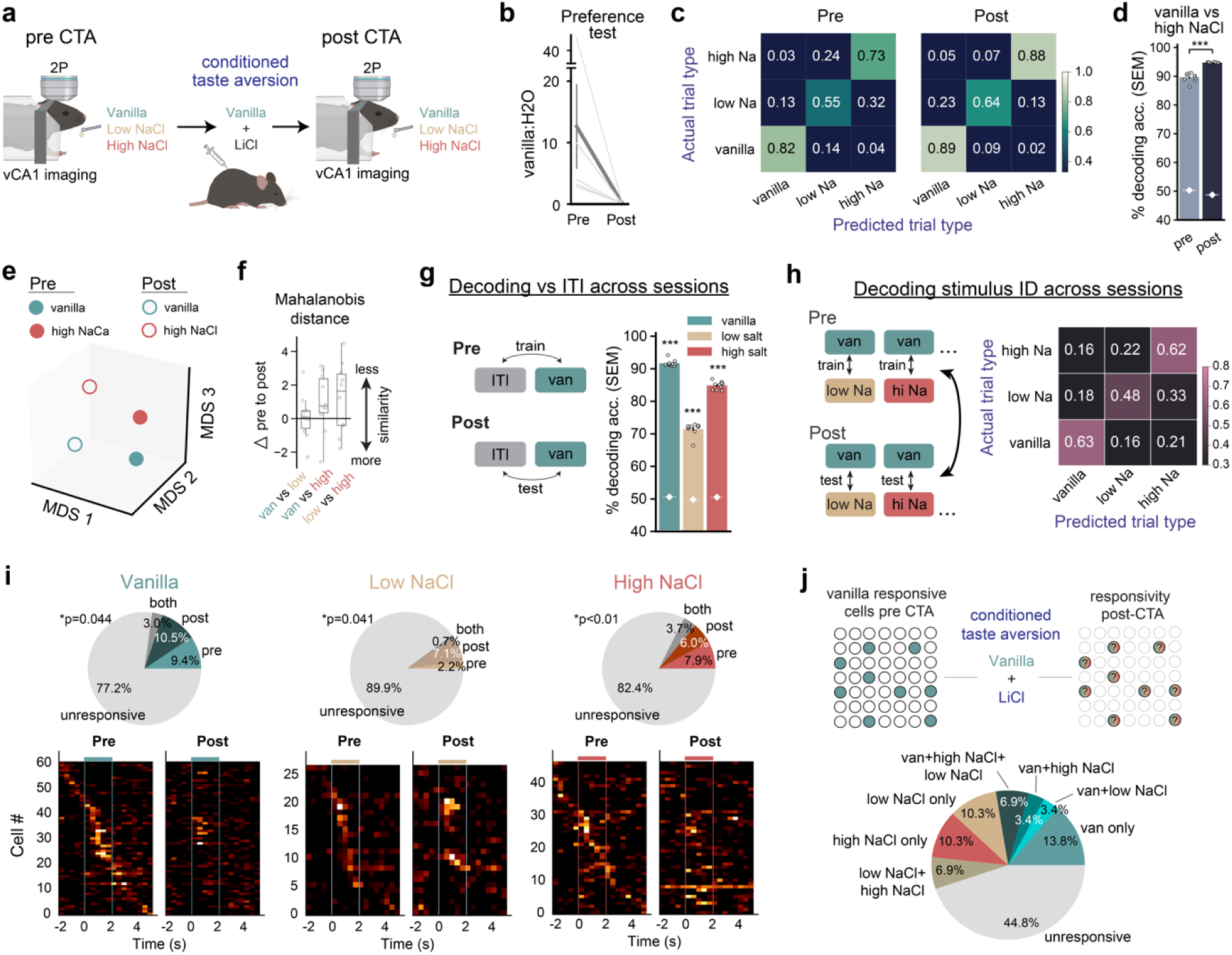
vCA1 Stimulus representations remain stable following reversal of stimulus valence via conditioned taste aversion (CTA). **a,** Task schematic. Before and after CTA administration, vCA1 was imaged via 2p microscopy while mice consumed liquids that differed in valence. Following exposure to an innately appetitive vanilla milk solution, mice were injected with lithium chloride to induce gastric malaise and subsequent aversion to vanilla milk. **b,** Prior to CTA, consumption of vanilla milk was highly preferred compared to water. Following CTA, mice avoided vanilla milk. **c,** Decoding confusion matrices comparing neural activity patterns during consumption of each liquid. In both pre- (left) and post-CTA (right) imaging sessions, decoding of stimulus identity (ascending diagonal) was most strongly represented by vCA1. **d,** Despite increased valence similarity between vanilla milk and high NaCl following CTA, neural representations of these stimuli become more distinct, as indicated by higher decoding accuracy in post vs pre. White diamonds and shading indicate mean +/− s.e.m. chance decoding accuracy. **e,** MDS visualization of vanilla and high NaCl stimulus representations, pre and post CTA. **f,** Mahalanobis distance analysis showed no increase in similarity between vanilla and high NaCl representations following CTA. **g,** Decoding vs baseline across sessions. Using pre-CTA recordings, a linear decoder was trained to discriminate neural activity occurring during intertrial interval (ITI) or stimulus consumption periods. ITI vs stimulus classification accuracy was then tested using post-CTA activity. Cross-session decoding accuracy was high for all stimuli indicating stability of stimulus representations across imaging sessions/CTA. **h,** Decoding trial type across sessions. A linear decoder trained to distinguish stimulus activity patterns prior to CTA, and testing using post-CTA activity was able to classify stimulus representations with high accuracy, further supporting stability of representations across CTA. **i,j,** Single-cell response properties. **i,** Top: pies showing percentage of neurons (n = 267) responsive to a particular stimulus during pre (lighter color), post (darker color), or across both sessions (dark grey). P-values reflect the observed proportion of cells responsive to across both sessions vs that expected by chance (see Methods). Bottom: Heatmaps plotting mean activity for all cells significantly responsive to a stimulus during pre and/or post sessions (pre heatmaps sorted by time of maximal activity, post sorted by pre). **j,** Top: analysis schematic. Cells responsive to vanilla during pre (n = 33) were tracked across sessions, and responsivity to each stimulus during post was measured. Bottom: pie chart of results. Vanilla-responsive neurons prior to CTA were equally likely to remap to low NaCl (appetitive) as high NaCl (aversive). \*\*\**P* < 0.001; \*\**P* < 0.01, \**P* < 0.05. All error bars indicate mean ± s.e.m. See Supplementary Table 1 for all statistical analysis details.

In both sessions, the neural activity evoked by each stimulus could be decoded from pre-stimulus background activity with high accuracy, indicating all three stimuli were salient to the animal (Supplementary Fig. 6A). When comparing the neural activity associated with each stimulus, decoding confusion matrices for pre- and post-CTA sessions indicated robust encoding of stimulus identity (Fig. 4**C**). Further, the accuracy of the stimulus identity decoding significantly increased from pre- to post-CTA sessions, consistent with the above and previously published (Biane et al., 2023) results showing that learning and repeated exposures lead to more discriminable representations (Supplementary Fig. 6B).

Of particular interest, however, was whether the vCA1 representation of vanilla milk, whose valence was inverted from appetitive to aversive following CTA, had come to more closely resemble the representation of the aversive high-NaCl solution. This was not the case. On the contrary, vanilla and high NaCl representations became more *dissimilar* following CTA. This was evidenced by significantly higher decoding accuracy of the two representations after CTA, as well as an increase in the Mahalanobis distance between the two representations (Fig. 4**D-F**). These results, once again, rebuff the view that stimulus valence is strongly encoded in vCA1, even after associative learning has drastically shifted the meaning an external stimulus holds for an animal.

If stimulus identity is robustly and stably encoded by vCA1, CTA administration should not appreciably alter the representation of vanilla milk, despite its change in valence. Indeed, we found this to be the case, as a decoder trained to differentiate stimulus-evoked activity from baseline activity during the pre-CTA session classified neural activity with high accuracy during post-CTA trials, and vice versa (Fig. 4**G**). Similarly, a decoder trained to differentiate neural activity evoked by all three stimulus-types during pre-CTA trials, performed well when classifying these stimuli during the post-CTA sessions, and vice versa. (Fig. 4**H**).

Single-cell results were consistent with these population level analyses, showing the proportion of co-responsive cells across pre- and post-CTA sessions for each stimulus was higher than expected by chance, signifying stability of stimulus representations (Fig. 4**I**). We also tracked cells responsive to vanilla milk during the pre-CTA session to see how they behaved after CTA (Fig. 4**J**), asking whether these cells would disproportionately remap and become responsive to high NaCl. This, however, did not occur. Initially vanilla-responsive neurons that switched responses were no more likely to remap to high NaCl than they were to low NaCl. Together, these data indicate stable vCA1 encoding of stimuli despite one of them having acquired a radically different meaning to the animal.

### vCA1 representations of emotionally neutral cues with acquired valence

Following cue-outcome associative learning, initially neutral, conditioned cues, such as a tone predictive of an upcoming shock, are capable of driving emotional responses (Anagnostaras et al., 2000; Gore et al., 2015). This suggests that learned cues, which initially have no inherent meaning, can become imbued with positive or negative valence. We therefore asked whether vCA1 representations might incorporate valence into representations of such stimuli once they have *learned* salience and valence.

To test this, mice were trained in a 4-odor associative learning paradigm, where one odor cue signaled the subsequent delivery of a weak shock (CS+_lsh_), one signaled a stronger shock (CS+_hsh_), one a sucrose reward (CS+_rew_) and one no outcome (CS-; Fig. 5**A**). Indicative of the animals having learned the cue-outcome associations, with training mice increased anticipatory licking behavior during the cue and trace period in sucrose trials, and reduced licking during this period in low and high shock trials (Fig. 5**B**).

**Figure 5.**
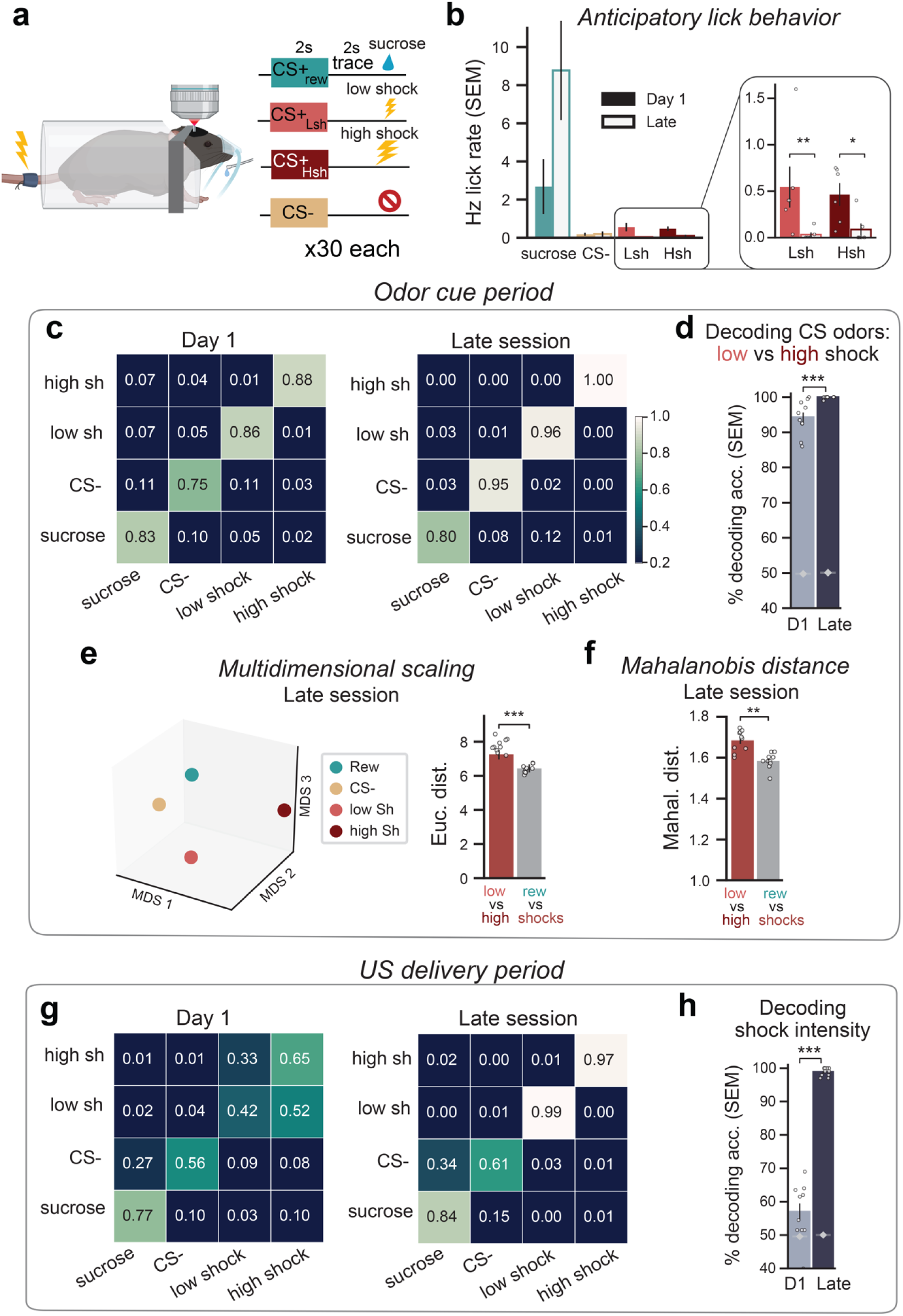
Stimulus intensity is encoded in vCA1, while learned cue valence is not. **a,** Schematic of trace associative learning paradigm. Four neutral odor cues were paired with one of four outcomes. Each trial type was presented 30 times, in pseudorandom order. **b,** Quantification of lick frequency that occurred after CS onset and before US. Anticipatory lick behavior increased for reward trials, while decreasing for both shock trial types, indicating animals had learned the cue-outcome associations. **c-f,** Analysis of the 2s odor delivery period to examine whether learned valence information is embedded in cue representations. **c,** Confusion matrices. True trial type on left, predicted trial type on bottom. Correct classification is along the ascending diagonal. classification accuracy is greater following training. **d,** Decoding specifically low vs high shock trials. Classification accuracy increases with training. White diamonds and shading indicate mean +/− s.e.m. chance decoding accuracy. **e,** MDS analysis shows the distance between low and high shock neural representations is significantly farther than between reward and shock representations. Thus, cue representations with learned similar valence do not cluster together. **f,** Mahalanobis distance analysis was consistent with MDS findings, showing greater distance between the negative valence shock trial types compared to the trial types with opposite valence (reward and shocks). **g,h,** Analysis during the 2s following US onset to examine whether stimulus intensity is encoded by vCA1. **g,** Same as in (**c**). Here US representations of low and high shock were discernible on the initial day of training, yet were often confused with one another when decoding, showing poor resolution for intensity coding. After training, however, the decoder made few errors. **h,** Decoding specifically low vs high shock trials. Classification accuracy is slightly above chance prior to learning and increases with training, indicating that distinct intensities of the same stimulus can be resolved via vCA1 activity, and increasingly so with training. \*\*\**P* < 0.001; \*\**P* < 0.01, \**P* < 0.05. All error bars indicate mean ± s.e.m. See Supplementary Table 1 for all statistical analysis details.

We reasoned that if valence is incorporated into the representation of an initially neutral cue that acquires predictive value to the animal, then the representations of the low and high shock odor cues should become more similar following cue-outcome learning. Instead, our results show that these representations became more separable, as decoding accuracy during the period of the odor cue presentation increased with learning (Fig. 5**C,D**). Furthermore, following training, the MDS geometric distance between low- and high-shock cue representations (same valence) was significantly greater than between the representations of the reward odor cue and shock cues (different valence; Fig. 5**E**). Again, this suggests that the learning process does not lead to valence information being incorporated into cue representations. Mahalanobis distance analyses corroborated these findings (Fig. 5**F**). Thus, analysis of odor cue activity shows an increase in discriminability with learning and argues against prominent valence coding in vCA1, not only for unconditioned stimuli, but learned, predictive cues as well.

### Representations of stimulus intensity in vCA1

Finally, we asked whether the different intensities of two stimuli of the same valence could be discriminated in vCA1 by measuring if vCA1 discriminated between the low and high shock amplitudes delivered in the above 4-odor associative learning task.

Early in training, analysis of vCA1 activity during the shock delivery periods revealed that the different intensity shocks could be decoded at better than chance. However, the classifier was prone to error, confusing representations of the two shock intensities with one another, albeit not with representations of sucrose delivery (Fig. 5**G,H**). With training, though, vCA1 improved discrimination of the two shock intensities, performing significantly better than early training periods and reaching a performance level of 90-100% accuracy. This suggests that the intensities of aversive stimuli are encoded by vCA1 and that discrimination between intensities improves with repeated exposure.

## Discussion

A fundamental function of the brain is to extract information from the environment and transform this into patterns of activity that support cognitive and emotional processing, which in turn promote appropriate behavioral selection. The vHPC plays a key role in this process and has been proposed to encode the emotional meaning of stimuli and contexts via generalized representations of positive or negative valence (Shpokayte et al., 2022; Turner et al., 2022; Xia and Kheirbek, 2020). Here, using a wide array of sensory inputs and experimental paradigms, we directly tested what information is embedded in the neural activity of vCA1.

These analyses have thoroughly shown that vCA1 neural activity encodes the identity, sensory modality and intensity of behaviorally salient stimuli. Further, these representations are relatively stable across days, but also accommodate improved neural discrimination of stimuli when they are presented multiple times. Unexpectedly, however, across multiple experimental designs and levels of analysis we failed to find convincing evidence that vCA1 contains representations of stimulus valence. Notably, even when an innately appetitive stimulus was conditioned to become aversive or when innately neutral odors were conditioned to be aversive or appetitive – processes that implicate the hippocampus (Aqrabawi and Kim, 2018; Koh et al., 2009; Li et al., 2017; Taxidis et al., 2020; Yun et al., 2023) – their representations in vCA1 were not imbued with a valence signal.

In well-studied scenarios that engage and depend on vCA1 function, subclasses of vCA1 neurons have been shown to encode distinct features of experience (AlSubaie et al., 2021; Ciocchi et al., 2015; Jimenez et al., 2020, 2018; Padilla-Coreano et al., 2016; Sánchez-Bellot et al., 2022; Tannenholz et al., 2014). For example, subpopulations of vCA1 neurons that project to the nucleus accumbens (NAc) encode rewarding experiences (Ciocchi et al., 2015; LeGates et al., 2018; Reed et al., 2018), while subpopulations of vCA1 neurons that are selectively active in anxiogenic environments tend to project to the medial prefrontal cortex and lateral hypothalamus (Adhikari et al., 2010; Jimenez et al., 2018; Padilla-Coreano et al., 2016). A prominent inference from these observations has been that vCA1 neurons directly encode and broadcast valence-related information of stimuli. Instead, we believe our results prompt a reassessment of vCA1 function, wherein the view that vCA1 classifies stimuli based on valence is replaced with models where vCA1 primarily classifies stimuli by their identity, and to which valence information is integrated via extended networks, such as the amygdala, where valence encoding is well documented (Beyeler et al., 2016; Felix-Ortiz et al., 2013; Gore et al., 2015; Zhang and Li, 2018). This main feature of vCA1 to encode individualized, highly separable stimulus representations, instead of a more general representation of valence, may serve to minimize generalization of emotional responses across similar stimuli and contexts, dysfunction of which is at the heart of myriad psychopathologies (Nees et al., 2015).

How do the current findings fit with past studies showing specific vHPC projection networks are biased toward processing positive or negative events? One possibility is that phenomena naturally associated with a particular valence are processed within designated pathways, such as cue-reward associations by vCA1 neurons projecting to NAc. In this scenario, valence is embedded within the projection pathway itself, while vCA1 neural activity transmits information related to stimulus identity, modality, intensity, etc. Another prospect is that there exists vCA1 subpopulations dedicated to processing any/all events of a particular valence. Here, valence information would be inherited from upstream inputs and passively transmitted by virtue of whether the recruited neurons are dedicated to processing positive or negative events; that is, valence information would be embedded within the *identity* of active neurons, and not their activity *patterns*. The mechanisms by which this may occur remain unclear, as vCA1 projection neurons receive broadly similar inputs, with a few modest exceptions (Gergues et al., 2020). Moreover, evidence suggests the valence associated with tagged ensembles of hippocampal neurons is fluid, as a specific HPC ensemble and its terminals can be manipulated to drive either approach or avoidance behavior (Redondo et al., 2014; Shpokayte et al., 2022). These latter studies suggest valence need not be fixed to specific pathways or cohorts of vHPC neurons but can be flexibly attached downstream.

Finally, our data suggest that vCA1 codes may be analogous to those found in dCA1, but operate on different incoming information. Just as dCA1 may integrate information about landmarks in the environment to generate a spatial map, vCA1 may integrate inputs related to motivation, emotional state, and the threatening or rewarding features of stimuli. In this manner, vCA1 may code for salient events and report their identity. Indeed, vCA1 engagement is highly dependent on the perceived salience of a stimulus (Biane et al., 2023). Thus, while vCA1 may not encode stimulus valence, *per se*, it may play a permissive role in selecting which stimuli, contexts and events warrant further emotional processing, and subsequently direct appropriate emotional and behavioral responses.

## Acknowledgements

The authors would like to thank Vijay Namboodiri, Loren Frank and Liam Drew for discussion and comments, and Sayi Boddu for technical assistance. JSB was supported by the Brain and Behavioral Research Foundation (NARSAD) and the Sandler PBBR Independent Postdoctoral Fellow Research Award. MAL was supported by NSF GR Fellowship. MAK was supported by NIMH (R01 MH108623, R01 MH111754, R01 MH117961), NIDCD (R01 DC019813) a One Mind Rising Star Award, a Research Grant from HFSP (Ref. No-RGY0072/2019), the Esther A. and Joseph Klingenstein Fund, the Pew Charitable Trusts, the McKnight Memory and Cognitive Disorders Award and The Ray and Dagmar Dolby Family Fund.

## Author contributions

Conceptualization: JSB, MAL, MAK

Methodology: JSB, MAL, MAK

Investigation: JSB, MAL, AF, HSC, LZZ, SH, DLAM, AOK, JXBP

Formal Analysis: JSB, MAL

Visualization: JSB, MAL, MAK

Funding acquisition: JSB, MAL, MAK

Supervision: JSB, MAK

Writing – original draft: JSB, MAK

Writing – review & editing: JSB, MAL, MAK

## Material and Correspondence

All datasets supporting the current study are available from the lead contact on request and will be publicly available from the Kheirbek lab github site (github.com/mkheirbek) prior to publication. Further information and requests for resources and reagents should be directed to and will be fulfilled by the Lead Contact, Mazen Kheirbek.

## Supplementary Materials

Supplementary Figs. 1 – 6

Supplementary Table 1

**Supplementary Figure 1.**
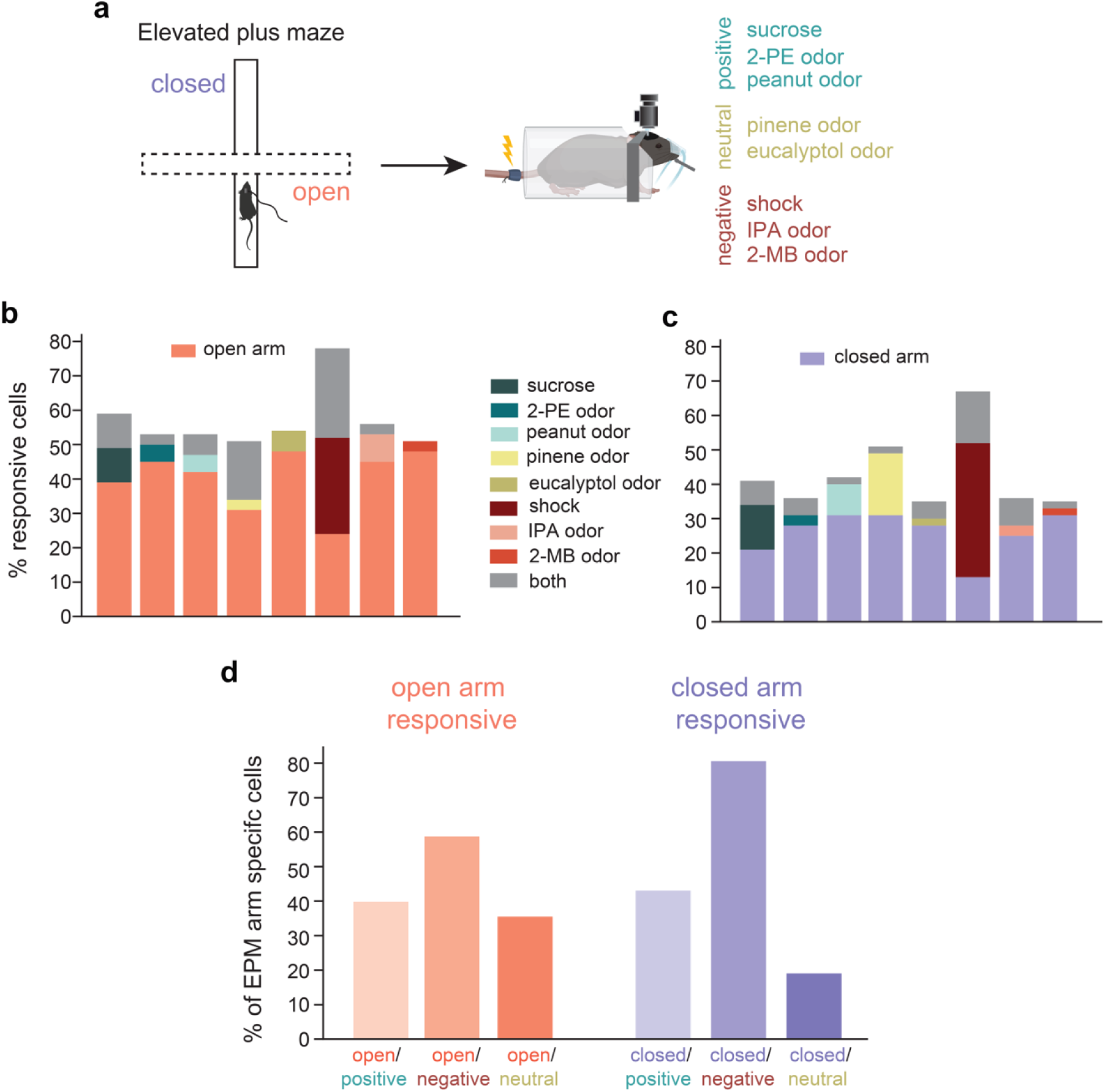
Cells responsive to open or closed arms of the EPM maze do not distinctly cluster with stimuli of a specific valence class. **a,** Experiment schematic. Neural activity was recorded via miniscope while animals freely explored an elevated plus maze (EPM). While recording from the same FOV, mice were subsequently exposed to a battery of stimuli while headfixed, which included odors, sucrose liquid and mild tail shocks. The valence categorization of each odor is shown on the right. **b,** Pairwise comparisons of open-arm responsive cells and every other stimulus presented. Percentage of cells responsive to both stimuli is shown in grey. **c,** Same as in (**b**), but for cells responsive to the closed arms**. d,** Percentage of open- or closed-arm responsive cells that were also responsive to any of the stimuli belonging to a specific valence class (i.e., the grey bars in (**b**) and (**d**), grouped by valence class). The presence of valence encoding would predict higher overlap for open-arm responsive cells (with open arm exploration being putatively anxiogenic) and the other negative stimuli. However, we found no consistent clustering of arm-selective cells with a specific valence class, as both open- and close-arm responsive cells showed appreciable overlap with each valence class.

**Supplementary Figure 2.**
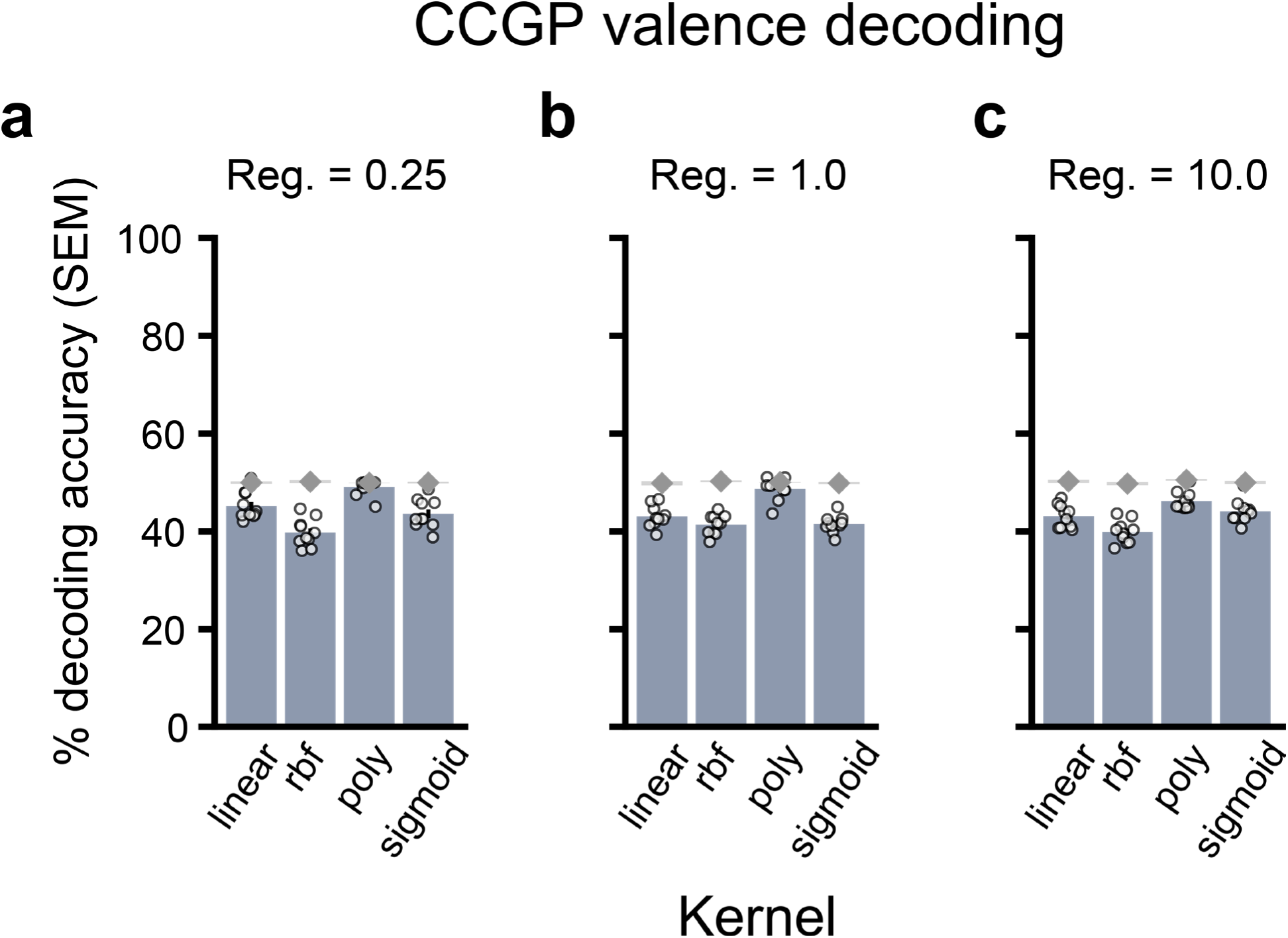
Valence decoding (sucrose/shock/female/predator) is no better than chance across a variety of kernel and regularization parameters. Valence decoding for results from Figure 1M using linear and nonlinear kernels at regularization values of (**a**) 0.25, (**b**) 1.0, or (**c**) 10.0. Decoding performance was no better than chance, regardless of parameters.

**Supplementary Figure 3.**
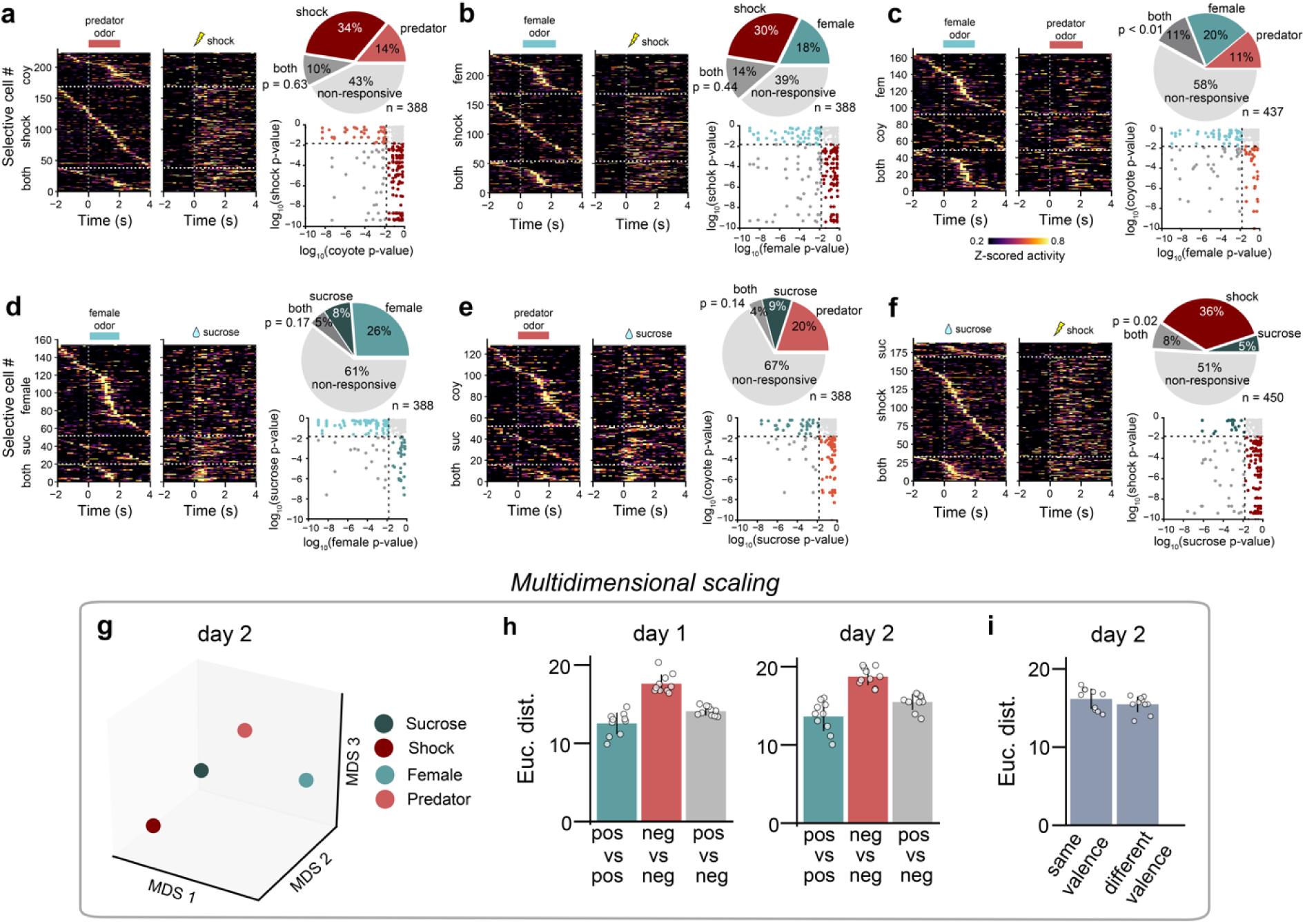
Day 2 responses to appetitive and aversive stimuli. **a-f,** As in Fig. 1d**-i** (day 1 data), but for day 2. Each panel shows the single-cell responses in a pairwise analysis of each stimulus pair. Heatmaps plot all cells significantly responsive to either of the two stimuli (stim 1 trials plotted on left, stim 2 trials on right). Pie charts show the percentage of cells that are responsive to each stimulus, both stimuli or none. **g,** MDS visualization, where the relationship between activity patterns is represented in geometrical space; the closer two points are in space, the more similar their activity patterns are. **h-i,** Clustering (lower Euclidean distance) between stimuli of a specific valence class vs across classes is indicative of valence encoding. However, distance between stimuli of opposing valence classes was no farther that between stimuli belonging to the same valence class, as seen in (**i**) for day 2, and Fig. 1n for day 1. All error bars indicate mean ± s.e.m. See Supplementary Table 1 for all statistical analysis details.

**Supplementary Figure 4.**
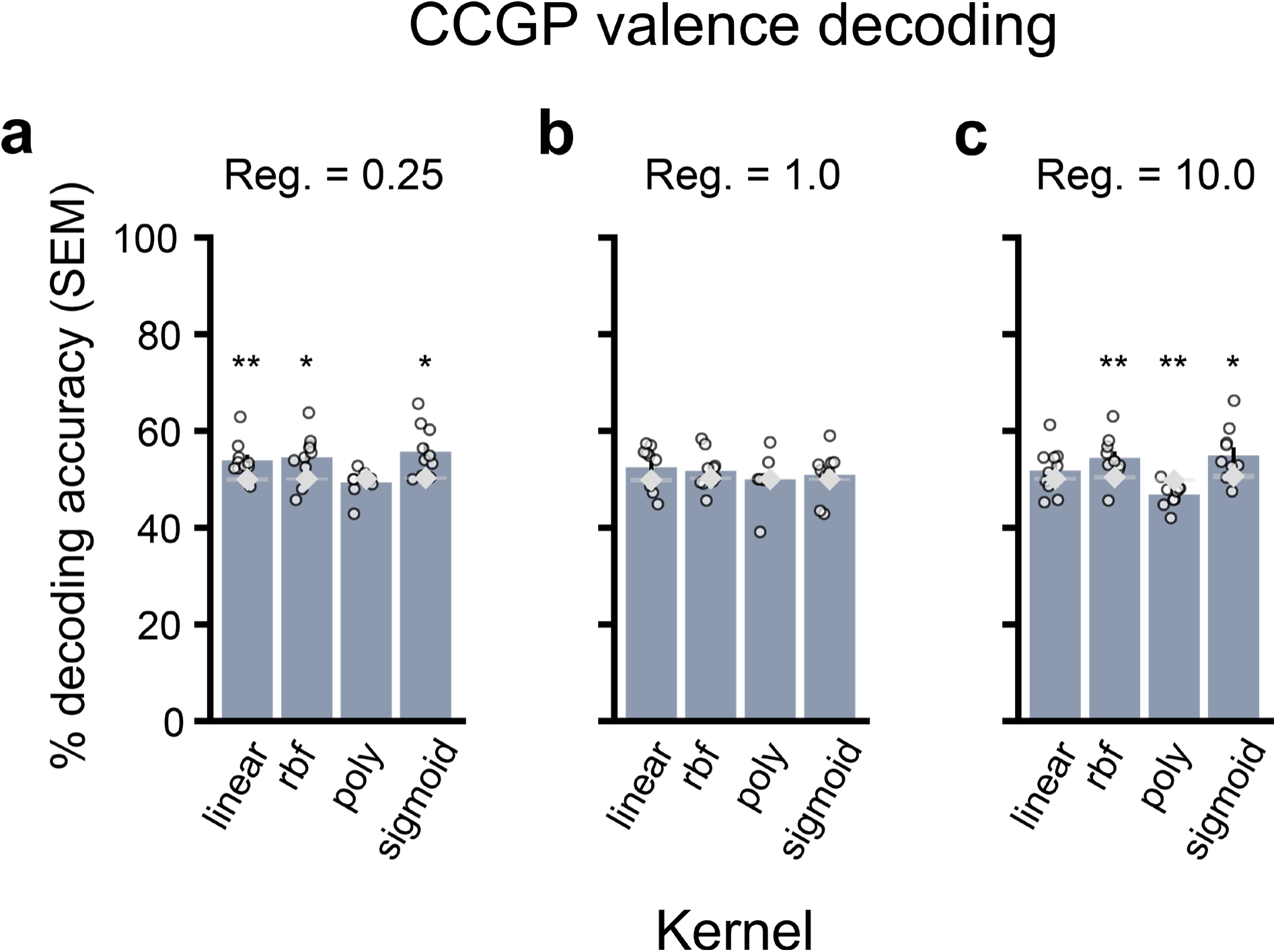
Valence decoding accuracy (sweet/umami/bitter/sour) across different kernel and regularization parameters. Valence decoding results from figure 3D (sweet/umami/bitter/sour) using linear and nonlinear kernels at regularization values of (**a**) 0.25, (**b**) 1.0, or (**c**) 10.0.

**Supplementary Figure 5.**
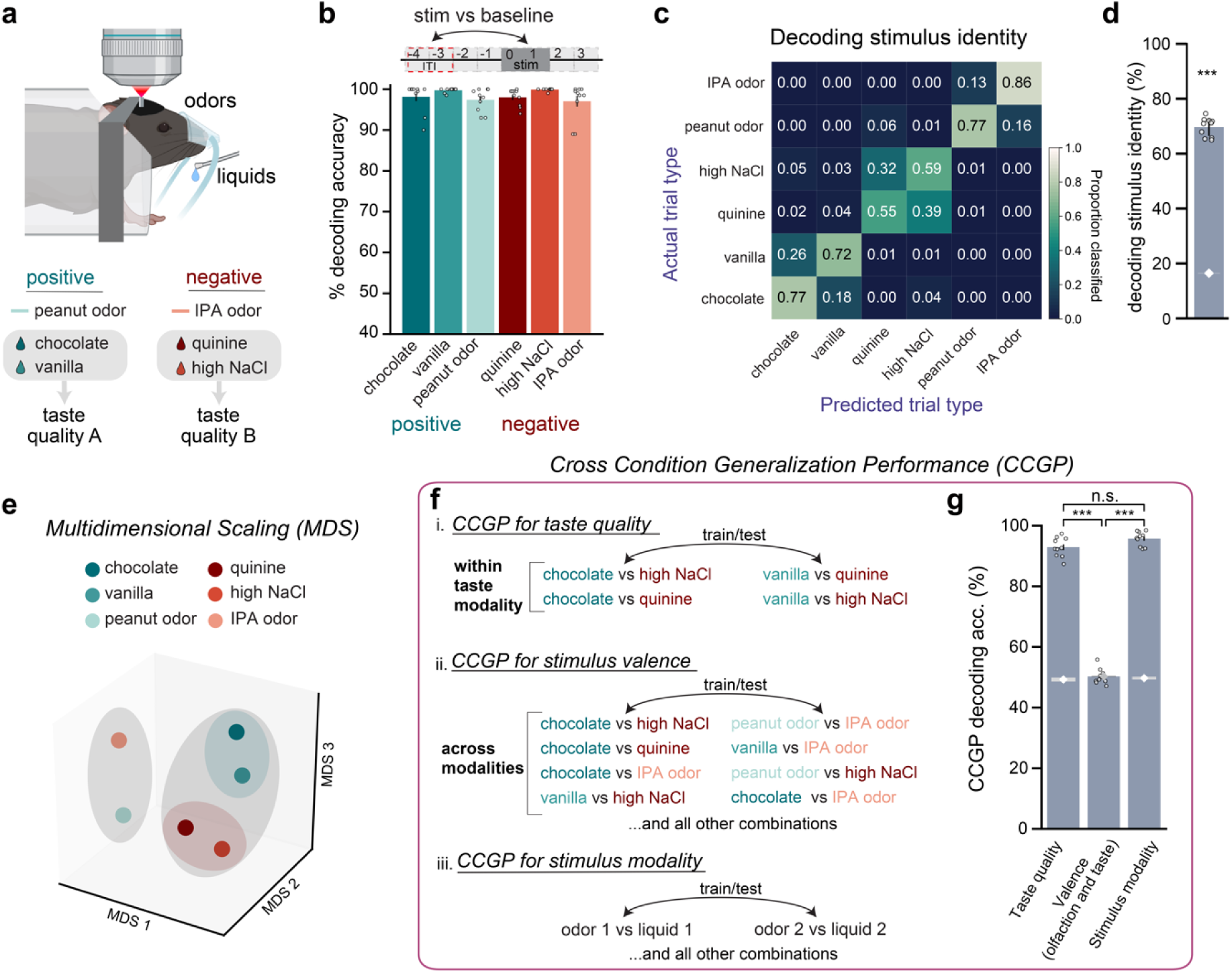
In vCA1, discrimination of stimulus identity and modality, generalization of taste quality. **a,** Experimental design. Mice were presented with 4 liquids (vanilla milk, chocolate milk, quinine or high NaCl) along with 2 odors, peanut oil and isopentylamine (IPA). **b,** The presence of each stimulus could be decoded from baseline (ITI) activity in vCA1. **c,d,** Decoding stimulus identity. White diamonds and shading indicate mean +/− s.e.m. chance decoding accuracy. **c**, Confusion matrix for all stimuli. While, in general, stimulus identity could be decoded well above chance. **d**, the confusion matrix shows the decoder made errors when discriminating between stimuli that activated similar taste pathways (vanilla and chocolate = sweet; quinine and high NaCl = bitter). **e,** MDS for visualizing the distance in representations shows a clustering of odor vs liquid representations (grey shading), as well as between sweet (teal shading) and bitter (red shading) taste qualities. **f,** Schematic for computing generalization performance of a decoder for taste quality (**i**), valence (**ii**), or stimulus modality (**iii**). **g,** The decoder successfully generalized across taste stimuli that were of the same taste quality but did not generalize across stimuli of similar valence. In addition, CCGP for stimulus modality performed significantly better than chance, consistent with our earlier results that modality but not valence is represented in vCA1. \*\*\**P* < 0.001; \*\**P* < 0.01, \**P* < 0.05. All error bars indicate mean ± s.e.m. See Supplementary Table 1 for all statistical analysis details.

**Supplementary Figure 6.**
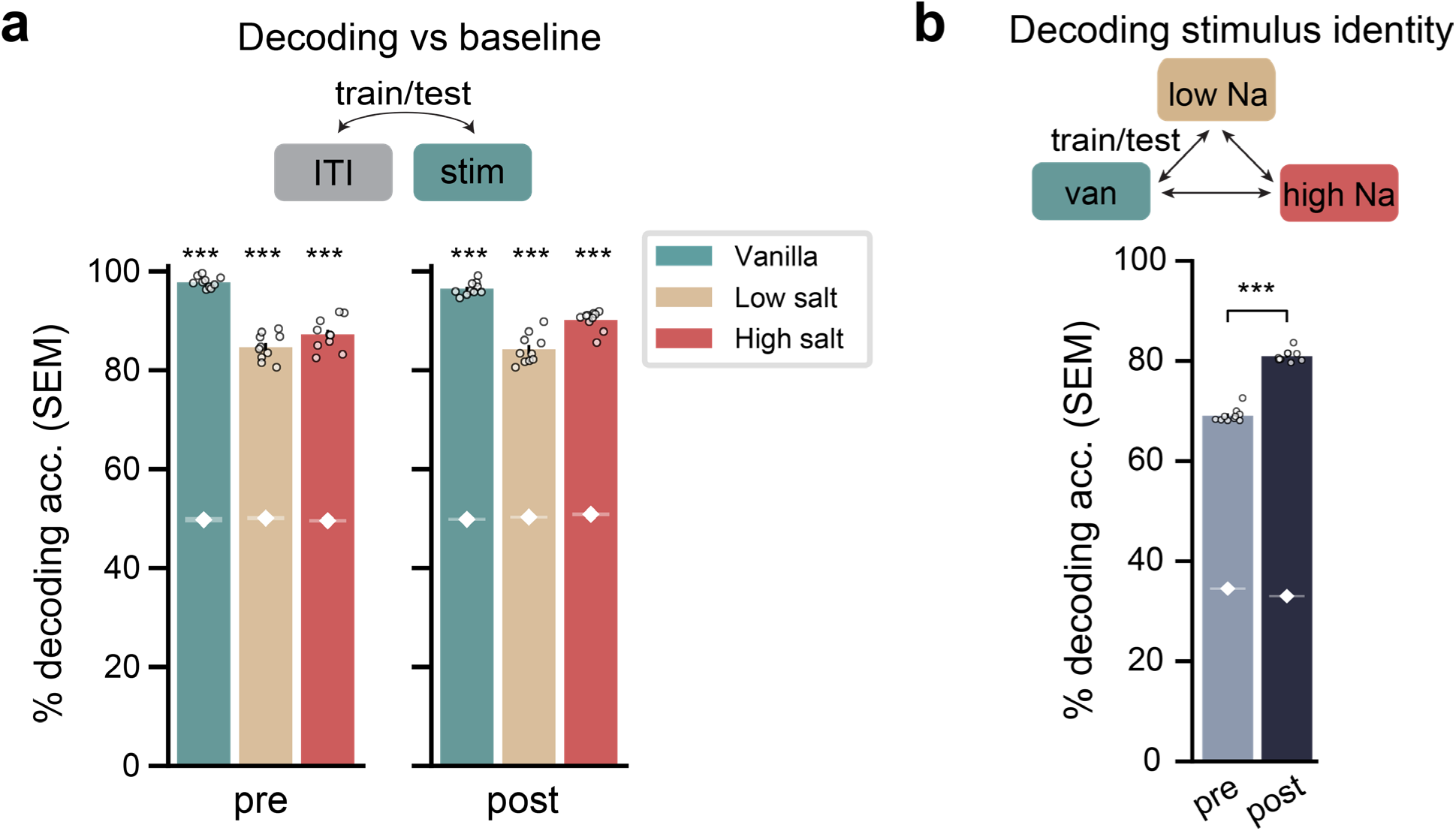
Liquid solutions are robustly represented by vCA1 before and after conditioned taste aversion, and become more separable with experience. **a,** A linear decoder could distinguish population activity during stimulus presentations versus during the intertrial interval (ITI) with high accuracy, for both pre- (left) and post-CTA (right) imaging sessions. White diamonds and shading indicate mean +/− s.e.m. chance decoding accuracy. **b,** Decoding of stimulus identity was significantly better in post compared to pre. \*\*\**P* < 0.001; \*\**P* < 0.01, \**P* < 0.05. All error bars indicate mean ± s.e.m. See Supplementary Table 1 for all statistical analysis details.

**Supplementary Table 1.** Statistical analysis parameters and results.

| Figure | Variable | Unit of Comparison | n | Test | Results |
| --- | --- | --- | --- | --- | --- |
| Fig 1D-I | overlap of stimulus selective cells (day 1) | cells | 410 cells (suc v fem; shock v pred; suc v pred; shock v fem); 474 cells (shock v suc); 454 cells (fem v pred) | p-value relative to shuffle (10,000) | suc v fem: $p = 0.11$<br>shock v pred: $p = 0.02$<br>shock v suc: $p = 0.02$<br>suc v pred: $p = 0.02$<br>fem v pred: $p < 0.01$<br>shock v fem: $p = 0.19$ |
| Fig 1L | decoding accuracy, stimulus identity vs chance | 10 decoding iterations | 400 cells from 7 mice | Mann-Whitney U | $p < 0.001$, effect-size ($r$) = 0.85 |
| Fig 1M | CCGP decoding accuracy, valence vs chance | 10 decoding iterations | 400 cells from 7 mice | Mann-Whitney U | $p = 0.34$ effect-size ($r$) = 0.22 |
| Fig 1N | CCGP decoding accuracy, stimulus modality vs chance | 10 decoding iterations | 400 cells from 7 mice | Mann-Whitney U | $p < 0.001$ effect-size ($r$) = 0.85 |
| Fig 1O | Euclidean distance between MDS coordinates (valence) | Euclidean distance | 400 cells from 7 mice | Mann-Whitney U | $p = 0.018$ effect-size ($r$) = 0.59 |
| Fig 1O | Euclidean distance between MDS coordinates (modality) | Euclidean distance | 400 cells from 7 mice | Mann-Whitney U | odor/non-odors: $p < 0.001$ effect-size ($r$) = 0.85<br>odors/odor vs non: $p < 0.001$ effect-size ($r$) = 0.85<br>non-odors/odor vs non: $p < 0.001$ effect-size ($r$) = 0.79 |
| Fig 2C-F | overlap of stimulus selective cells across days | cells | 325 cells (suc; shock); 296 cells (fem; pred) | p-value relative to shuffle (10,000) | suc: $p = 0.025$<br>shock: $p = 0.002$<br>fem: $p = 0.033$<br>pred: $p < 0.001$ |
| Fig 2H | decoding accuracy, stimulus identity day 1 vs day 2 | 10 decoding iterations | d1 = 400 cells from 7 mice<br>d2 = 388 cells from 7 mice | Mann-Whitney U | $p < 0.001$ effect-size ($r$) = 0.85 |
| Fig 2I | CCGP decoding accuracy, valence day 1 vs day 2 | 10 decoding iterations | d1 = 400 cells from 7 mice<br>d2 = 388 cells from 7 mice | Mann-Whitney U | $p = 0.09$ effect-size ($r$) = 0.46 |
| Fig 3B | decoding accuracy, stimulus vs ITI baseline period (vs chance) | 10 decoding iterations | 967 cells from 7 mice | Mann-Whitney U | sweet: $p < 0.001$ effect-size ($r$) = 0.85<br>umami: $p < 0.001$ effect-size ($r$) = 0.85 |
| | | | | | bitter: $p < 0.001$<br>effect-size ($r$) = 0.85<br>sour: $p < 0.001$<br>effect-size ($r$) = 0.85 |
| Fig 3D | decoding accuracy, stimulus identity vs chance | 10 decoding iterations | 967 cells from 7 mice | Mann-Whitney U | $p < 0.001$, effect-size ($r$) = 0.85 |
| Fig 3F | Euclidean distance between MDS coordinates (valence) | Euclidean distance | 967 cells from 7 mice | Mann-Whitney U, Bonferroni correction for $n=2$ | +/+ vs -/-: $p = 1.0$, effect-size ($r$) = 0.14<br>'+/+ vs +/-: $p = 0.61$, effect-size ($r$) = 0.24<br>'-/- vs +/-: $p = 1.0$, effect-size ($r$) = 0.16 |
| Fig 3G | Mahalanobis distance between stimulus representations (valence) | Mahalanobis distance | 967 cells from 7 mice | Mann-Whitney U, Bonferroni correction for $n=2$ | +/+ vs -/-: $p = 0.48$, effect-size ($r$) = 0.27<br>'+/+ vs +/-: $p = 1.0$, effect-size ($r$) = 0.15<br>'-/- vs +/-: $p = 0.24$, effect-size ($r$) = 0.35 |
| Fig 3H | CCGP decoding accuracy, stimulus valence vs chance | 10 decoding iterations | 967 cells from 7 mice | Mann-Whitney U | $p = 0.002$, effect-size ($r$) = 0.69 |
| Fig 4B | liquid preference: vanilla:water delivery ratio | session (pre CTA vs post CTA) | 5 mice | Mann-Whitney U | $p = 0.012$, effect-size ($r$) = 0.88 |
| Fig 4D | pre vs post decoding accuracy, vanilla milk vs high NaCl | 10 decoding iterations | 267 cells from 5 mice (cells registered across sessions) | Mann-Whitney U | $p < 0.001$, effect-size ($r$) = 0.85 |
| Fig 4G | across-session decoding accuracy, stimulus vs ITI baseline period | 10 decoding iterations | 267 cells from 5 mice (cells registered across sessions) | Mann-Whitney U | vanilla: $p < 0.001$, effect-size ($r$) = 0.85<br>low NaCl: $p < 0.001$, effect-size ($r$) = 0.85<br>high NaCl: $p < 0.001$, effect-size ($r$) = 0.85 |
| Fig 4I | pre and post overlap of stimulus selective cells | cells | 267 cells from 5 mice (cells registered across sessions) | p-value relative to shuffle (10,000) | vanilla: $p=0.43$<br>low NaCl: $p = 0.04$<br>high NaCl: $p < 0.001$ |
| Fig 5B | mean lick rate (Hz) | session (day 1 vs late) | 6 mice | Mann-Whitney U | low shock: $p = 0.009$ , effect- size (r) = 0.76<br>high shock: p = 0.028, effect-size (r) = 0.65 |
| Fig 5D | decoding odor cue accuracy across sessions, low shock trials vs high shock trials | 10 decoding iterations | d1 = 141 cells from 6 mice<br>late = 133 cells from 6 mice | Mann-Whitney U | p < 0.001, effect-size (r) = 0.85 |
| Fig 5E | Euclidean distance between MDS odor cue coordinates | Euclidean distance | d1 = 141 cells from 6 mice<br>late = 133 cells from 6 mice | Mann-Whitney U | p = 0.018<br>effect-size (r) = 0.59 |
| Fig 5F | Mahalanobis distance between odor cue representations | Mahalanobis distance | d1 = 141 cells from 6 mice<br>late = 133 cells from 6 mice | Mann-Whitney U | p = 0.001<br>effect-size (r) = 0.74 |
| Fig 5H | decoding odor cue accuracy across sessions, low shock trials vs high shock trials | 10 decoding iterations | d1 = 141 cells from 6 mice<br>late = 133 cells from 6 mice | Mann-Whitney U | p < 0.001, effect-size (r) = 0.85 |
| Fig S3A-F | overlap of stimulus selective cells (day 2) | cells | 388 cells (suc v fem; shock v pred; suc v pred; shock v fem); 450 cells (shock v suc); 437 cells (fem v pred) | p-value relative to shuffle (10,000) | suc v fem: p = 0.17<br>shock v pred: p = 0.63<br>shock v suc: p = 0.02<br>suc v pred: p = 0.14<br>fem v pred: p < 0.01<br>shock v fem: p = 0.44 |
| Fig S3H | Euclidean distance between MDS stimulus coordinates day 1 | Euclidean distance pos v pos VS neg v neg | 401 cells from 7 mice | Mann-Whitney U, Bonferroni correction for n=2 | p < 0.001<br>effect-size (r) = 0.85 |
| Fig S3H | Euclidean distance between MDS stimulus coordinates day 1 | Euclidean distance pos v pos VS pos v neg | 401 cells from 7 mice | Mann-Whitney U, Bonferroni correction for n=2 | p < 0.001<br>effect-size (r) = 0.85 |
| Fig S3H | Euclidean distance between MDS stimulus coordinates day 1 | Euclidean distance pos v neg VS neg v neg | 401 cells from 7 mice | Mann-Whitney U, Bonferroni correction for n=2 | p = 0.009<br>effect-size (r) = 0.64 |
| Fig S3H | Euclidean distance between MDS stimulus coordinates day 2 | Euclidean distance pos v pos VS neg v neg | 388 cells from 7 mice | Mann-Whitney U, Bonferroni correction for n=2 | p < 0.001<br>effect-size (r) = 0.85 |
| Fig S3H | Euclidean distance between MDS stimulus coordinates day 2 | Euclidean distance pos v pos VS pos v neg | 388 cells from 7 mice | Mann-Whitney U, Bonferroni correction for n=2 | p < 0.001<br>effect-size (r) = 0.85 |
| Fig S3H | Euclidean distance between MDS stimulus coordinates day 2 | Euclidean distance pos v neg VS neg v neg | 388 cells from 7 mice | Mann-Whitney U, Bonferroni correction for n=2 | p = 0.051<br>effect-size (r) = 0.51 |
| Fig S3I | Euclidean distance between MDS stimulus coordinates (day 2) | Euclidean distance | 388 cells from 7 mice | Mann-Whitney U | p = 0.37<br>effect-size (r) = 0.3 |
| Fig S5B | decoding accuracy, stimulus vs ITI baseline period (vs chance) | 10 decoding iterations | 123 cells from 4 mice | Mann-Whitney U | choc: p < 0.001, r = 0.85<br>van: p < 0.001, r |
| | | | | | = 0.85<br>pnut: $p < 0.001$,<br>$r = 0.85$<br>qui: $p < 0.001$, $r = 0.85$<br>NaCl: $p < 0.001$,<br>$r = 0.85$<br>IPA: $p < 0.001$, $r = 0.85$ |
| Fig S5D | decoding accuracy, stimulus identity vs chance | 10 decoding iterations | 123 cells from 4 mice | Mann-Whitney U | $p < 0.001$, effect-size ($r$) = 0.85 |
| Fig S5G | decoding accuracy, stimulus vs ITI baseline period (vs chance) | 10 decoding iterations | 123 cells from 4 mice | Mann-Whitney U | taste quality: $p < 0.001$, effect-size ($r$) = 0.85<br>valence: $p = 0.47$, effect-size ($r$) = 0.17<br>modality: $p < 0.001$, effect-size ($r$) = 0.85 |
| Fig S5G | decoding accuracy, taste quality vs valence (olfaction and taste) | 10 decoding iterations | 123 cells from 4 mice | Mann-Whitney U, Bonferroni correction for $n=2$ | $p < 0.001$, effect-size ($r$) = 0.85 |
| Fig S5G | decoding accuracy, taste quality vs stimulus modality | 10 decoding iterations | 123 cells from 4 mice | Mann-Whitney U, Bonferroni correction for $n=2$ | $p = 0.24$, effect-size ($r$) = 0.35 |
| Fig S5G | decoding accuracy, valence (olfaction and taste) vs stimulus modality | 10 decoding iterations | 123 cells from 4 mice | Mann-Whitney U, Bonferroni correction for $n=2$ | $p = 0.08$, effect-size ($r$) = 0.47 |
| Fig S6A | Pre session decoding accuracy, stimulus vs ITI baseline period (vs chance) | 10 decoding iterations | 267 cells from 5 mice (cells registered across sessions) | Mann-Whitney U | vanilla: $p < 0.001$, $r = 0.85$<br>low NaCl: $p < 0.001$, $r = 0.85$<br>high NaCl: $p < 0.001$, $r = 0.85$ |
| Fig S6A | Post session decoding accuracy, stimulus vs ITI baseline period (vs chance) | 10 decoding iterations | 267 cells from 5 mice (cells registered across sessions) | Mann-Whitney U | vanilla: $p < 0.001$, $r = 0.85$<br>low NaCl: $p < 0.001$, $r = 0.85$<br>high NaCl: $p < 0.001$, $r = 0.85$ |
| Fig S6B | pre vs post decoding accuracy, stimulus identity vs chance | 10 decoding iterations | 267 cells from 5 mice (cells registered across sessions) | Mann-Whitney U | $p < 0.001$, effect-size ($r$) = 0.85 |

## Methods

### KEY RESOURCES TABLE

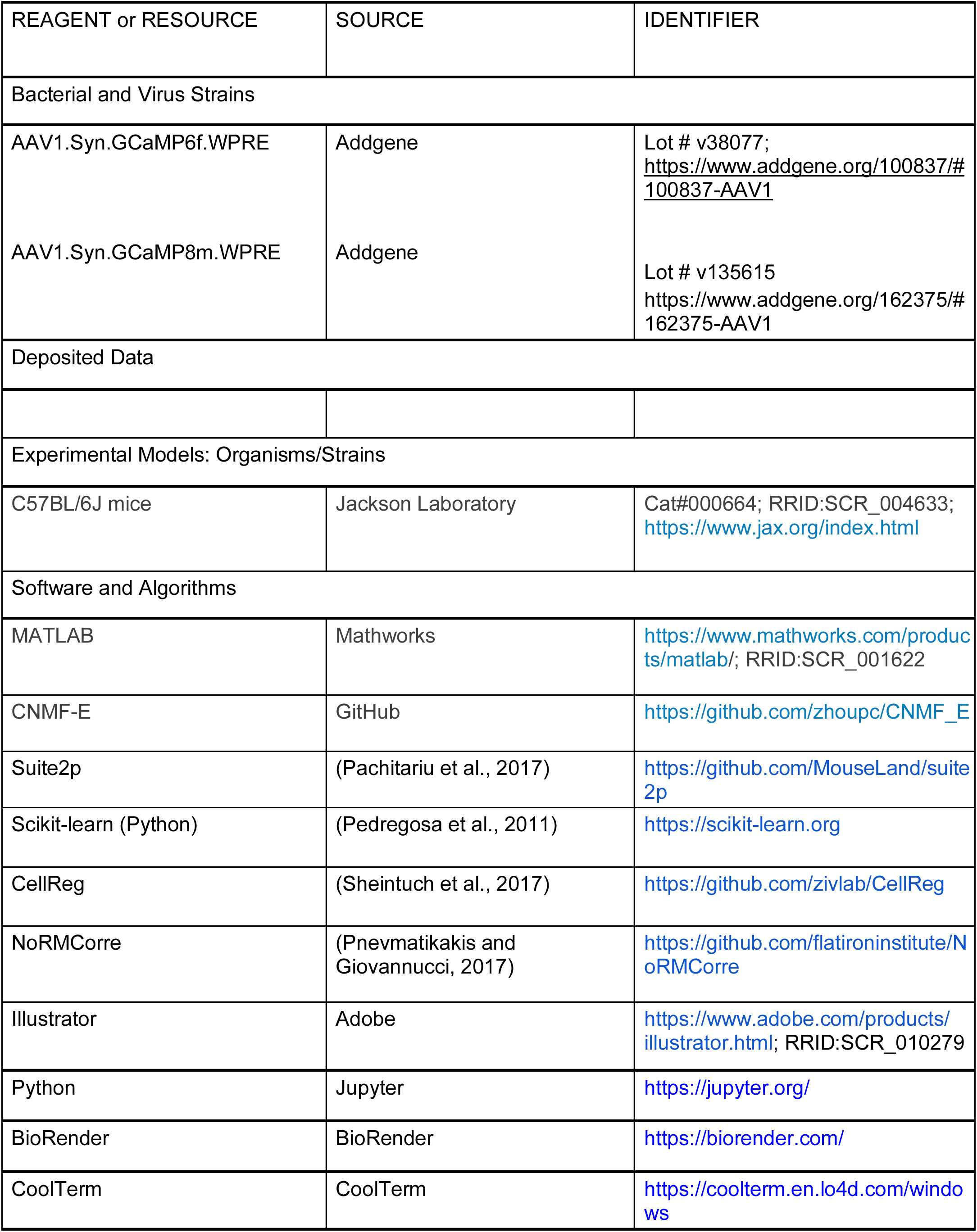
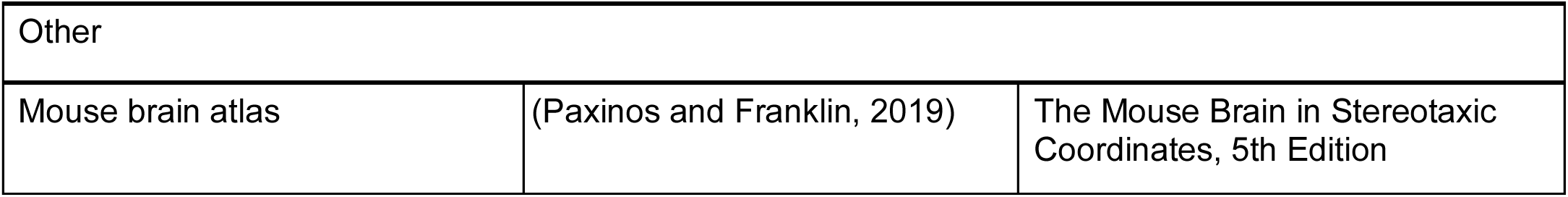

### Contact for reagents and resource sharing

Further information and requests for resources and reagents should be directed to and will be fulfilled by the Lead Contact, Mazen Kheirbek.

### Materials Availability

This study did not generate new unique reagents.

### Data and Code Availability

The datasets and analysis code supporting the current study are available from the lead contact on request.

### Mice

All procedures were conducted in accordance with the U.S. National Institutes of Health Guide for the Care and Use of Laboratory Animals and the institutional Animal Care and Use Committees at University of California, San Francisco (UCSF). Adult male and female C57BL/6J mice were supplied by Jackson Laboratory. Mice were co-housed with littermates (2–5 per cage) in a temperature (22–24 °C) and humidity (40–60%) controlled environment on a 12-h light/dark cycle, with experiments performed during the light phase. All mice were randomly assigned to experimental conditions. Data collection and analysis were not performed blind to the conditions of the experiments. All experiments included both male and female mice, with the exception of experiment 1, which only included sexually naïve male mice so that female urine odor would act as a highly appetitive stimulus.

### Surgery

Animals were 11 – 15 weeks old at time of surgery. Mice were anesthetized with 1.5% isoflurane with an O2 flow rate of 1 L / min, and head-fixed in a stereotactic frame (David Kopf, Tujunga, CA). Eyes were lubricated with an ophthalmic ointment, and body temperature was maintained at 37°C with a warm water recirculator (Stryker, Kalamazoo, MI). The fur was shaved and incision site sterilized prior to beginning surgical procedures. Lidocaine, meloxicam, and slow-release buprenorphine were provided for analgesia.

GCaMP8m (experiments 3, 4 and 5) or GCaMP6f (experiments 1, 2 and 6) virus injection and GRIN lens implantation were conducted using methods previously described (Jimenez et al., 2018). Briefly, a craniotomy was made over the lens implantation site and dura was removed from the brain surface and cleaned with sterile saline and absorptive spears (Fine Science Tools, Foster City, CA). A nanoject syringe (Drummond Scientific, Broomall, PA) was used to deliver GCaMP to vCA1 (left hemisphere). vCA1 coordinates were −3.16 A/P and −3.25 M/L. 150nl of virus was injected at each depth of −3.85, −3.55 and −3.3 (450nl total volume) with respect to bottom of skull at the medial edge of the craniotomy. The needle was held in place for > 5 minutes prior to moving to the next D/V coordinate and remained in place for 10 minutes following the final injection before slowly removing from the brain. AAV1-SYN-GCaMP6f-WPRE-Sv40 (titer: 1.97E+13) was supplied from the University of Pennsylvania viral vector core and diluted 1:3 in 1x sterile PBS before injections. AAV1-SYN-jGCaMP8m-WPRE (titer: 2.4E+13) was supplied from Addgene and diluted 1:3 in 1x sterile PBS before injections. Following virus injection, a 0.6mm diameter GRIN lens (Inscopix, Palo Alto, CA) was slowly lowered in 0.1 mm D/V steps and then fixed to the skull with Metabond adhesive cement (Parkell, Edgewood, NY). vCA1 lens coordinates were −3.16 A/P, −3.5 M/L and −3.5 D/V (from bottom of skull at craniotomy). A custom-made titanium headbar was then attached to the skull using dental cement (Dentsply Sinora, Philadelphia, PA). A baseplate and cover (Inscopix, Palo Alto, CA) was also cemented on to protect the lens. In total, 27 animals were imaged.

### Verification of imaging sites and histological analysis

Ventral CA1 imaging sites were verified in each animal included in final analysis. After all imaging sessions were completed, mice were injected with a lethal dose 2:1 ketamine/xylazine solution intraperitoneally. With the heart still beating, mice were perfused transcardially using 4% PFA solution. Brains were extracted and placed in 4% PFA solution for 2-3 days to allow further fixation. After saturating with a 30% sucrose solution, coronal slices of 50-micron width were collected using a Leica SM2000 microtome. Slices were collected in 1x PBS solution and mounted onto glass slides, coverslipped with Fluoromount G with DAPI (Southern Biotech, Birmingham, AL).

### Experimental paradigms

Animals used for experiments, one, four and six had been trained on a headfixed, cue-outcome trace conditioning paradigm prior. Multisession paradigms consisted of one session/day, occurring at roughly the same time each day.

#### Experiment 1: Sucrose, shock, predator urine, female conspecific urine

Only male mice were used for this experiment, due to female conspecific urine used as a positive stimulus. In all other experiments, both male and female mice were used. Animals were water restricted to ∼85-90% ad lib weight. Mice were headfixed, and a nose cone placed over their snout through which individual odors were delivered at a flow rate of ∼1L per minute (200A olfactometer, Aurora Scientific, Aurora, ON, Canada). Each odor presentation lasted 2 sec. Odors were delivered to mice via a customized nose cone, which contained an outlet where a gentle vacuum was applied to evacuate residual odor. Additionally, an ongoing charcoal filter vacuum system (Hydrobuilders Inc.) was used to evacuate any residual odors. Odors consisted of an attractive female odor (800 µL mineral oil and 300 µL female urine pooled from several (3+) females of 3-6 months old, collected fresh daily) and an aversive coyote odor (800 µL mineral oil, 300 µL coyote urine, Maine Outdoor Solutions; Hermon, Maine). Mild electric shocks (0.3 mA, 500 ms) were delivered via a custom-made tail cuff driven by a precision animal shocker (Coulbourn Instruments, Holliston, MA). Liquid sucrose solution (10% sucrose, 0.03% NaCl in water, 3 µL per presentation) was delivered via a solenoid-gated gravity feed into a spout set in front of the mouth. Contact with a lick spout positioned in front of the mouth was measured using a capacitive touch MPR121 sensor (SparkFun, Boulder, CO). Stimulus delivery and sensor reading was controlled by an Arduino Mega with custom circuit boards (adapted from OpenMaze.org) and recorded via CoolTerm software. Mice were exposed to the following experimental paradigm for two consecutive days: 30 presentations of odors (15 pseudorandom alternating presentations of coyote urine and female urine, 30 presentations of sucrose rewards (2µl each), and 10 presentations of tail shocks. For every trial, the interval between successive stimulus presentations was selected from a uniform distribution spanning 15-30 sec. Imaging was briefly halted between odor and sucrose/shock epochs in order to position the lick spout and tail shock cuff. Because imaging was not continuous, cells imaged during odor and sucrose/shock sessions required registration across these epochs. Therefore, the total number of cells differ when comparing stimuli within the same recording session vs across sessions, where a subset of cells could not be reliably registered across sessions. A total of 7 distinct FOVs from 6 mice were included in the data set.

#### Experiment 2: EPM + stimulus battery

*Elevated Plus Maze*. Mice were placed in a standard EPM sized maze (13.5’’ height of maze from floor, 25’’ full length of each armtype, 2’’ arm width, 7’’ tall closed arms, with 0.5’’ tall/wide ledges on the open arms made by Maze Engineers, Skokie, IL), with 450 light lux centered over the open arms to promote avoidance. Mice were placed in the center region of the maze and were allowed to explore for 20 minutes while recording behavior with a webcam EthoVision XT 10 (Noldus, Leesburg, VA). Behavior was tracked using DeepLabCut software (version 2.0.6.2) (Mathis et al., 2018; Nath et al., 2019) and custom written python code (https://www.google.com/url?q=https://www.nature.com/articles/s41593-018-0209-y&sa=D&source=docs&ust=1694216523679815&usg=AOvVaw0YAy7kvZKam_0HeOZdHHk4). We tracked head, body, and tail positions (nose, start of tail, end of tail), and used the body position to determine open, center and closed arm occupancy. Specifically, we labeled 40 number of frames taken from 5 videos/animals, then 95% was used for training. We used a ResNet-50-based neural network with default parameters for (1003000 x) 2 training iterations. We used a p-cutoff of .9 to condition the X, Y coordinates for future analysis after validation then used custom code to further filter data for discontinuities. This network was then used to analyze videos from similar experimental settings.

*Odors and Sucrose and Shock*. In two separate experimental contexts, head-fixed mice were exposed to either a panel of valenced odors or sucrose liquid and shocks while being imaged. Sucrose and shock were given as previously described, except 60 rewards and 30 shocks were given with all trials being used for analysis. Six odors, including two appetitive (peanut oil, 2-phenylethanol), two aversive (isopentylamine, 4-methylthiazole), and two neutral (alpha-pinene, eucalyptol), were presented 25 times each. Odors were diluted to 10% in 10 ml total volume mineral oil. The inter-trial interval between subsequent stimuli was chosen as a random sample from a uniform distribution between 12 and 18 seconds. Mice were headfixed, and a nose cone placed over their snout through which individual odors were delivered at a flow rate of ∼1L per minute. Each odor presentation lasted 2 sec.

#### Experiment 3: Liquids

Four liquids were used, each known to activate specific taste representations (Chen et al., 2011). Sweet: 20% sucrose. Umami: 70 mM MPG (monopotassium glutamate) plus 1 mM IMP (inosine 5′-monophosphate). Bitter: 20uM cycloheximide. Sour: 2.5% citric acid. All solutions were diluted in deionized water. Stimuli were interleaved and pseudorandomly presented. Each stimulus was presented 12 times. The inter-trial interval between subsequent stimuli was chosen as a random sample from a uniform distribution between 15 and 25 seconds. For liquid delivery, one of four delivery spouts was rotated into place directly in front of the animal at the onset of the appropriate trial. Each spout delivered a single liquid only. All lick spouts remained out of reach of the animal during the ITI. The appropriate lick spout was rotated into place at trial onset (∼300ms), followed by liquid delivery ∼150ms after rotation was completed. The lick spout was subsequently rotated back to home position 3 sec post liquid delivery. Mouse behavior was manually inspected during the task and trials where a liquid was not consumed within 1 sec of delivery were excluded from further analysis (9.5% of trials excluded).

Animals were water restricted to ∼90% ad lib weight at time of recording. A total of 14 distinct FOVs from 7 mice were included in the data set.

#### Experiment 4: Tastants and odors

A total of 6 odors (peanut butter/peanut oil mixture 100%, 2-phenylethanol 10%, isopentylamine 30%, 4-methylthiazole 30%, valeraldehyde 1.5%, pentanedione 10%) and 4 liquids (Nesquik chocolate milk diluted 50% in water, Nesquik vanilla milk diluted 50% in water, 10mM quinine, 550mM NaCl) were used. Odors were diluted to 10 ml total volume in mineral oil. Stimuli were interleaved and pseudorandomly presented. Each stimulus was presented 11 times, with the first 10 trials included for analysis. The inter-trial interval between subsequent stimuli was chosen as a random sample from a uniform distribution between 15 and 25 seconds. Mice were headfixed, and a nose cone placed over their snout through which individual odors were delivered at a flow rate of ∼1L per minute. Each odor presentation lasted 2 sec. Liquid delivery followed the same methodology as experiment 3. Video recordings of mouse behavior were manually inspected and used to exclude trials where a liquid was not consumed within 1 sec of delivery (6.6% of trials excluded).

Animals were water restricted to ∼90% ad lib weight at time of recording. A total of 8 distinct FOVs from 4 mice were included in the data set.

#### Experiment 5: Conditioned taste aversion (CTA)

Mice were first habituated to the liquid delivery chamber for 4 days, with water freely available. The morning of day 5, mice were given access to water to minimize any general rewarding properties of hydration across the distinct solutions provided during imaging. Four hours later, mice were imaged via 2-photon microscopy while headfixed and receiving liquids. Three liquids were used: vanilla milk (Nesquik, 50%), low NaCl solution (125mM) or high NaCl solution (1.1M). All solutions were diluted in deionized water. During imaging, liquids were delivered as in experiments 3 and 4. The following day (day 6), liquid preference was tested. Mice were placed in the liquid delivery chamber for 30 minutes and given free access to vanilla milk and water, each delivered from a separate spout. A 3uL liquid delivery was triggered by a lick to the corresponding spout and was controlled by an Arduino microcontroller. Liquid deliveries were recorded via CoolTerm software. Five minutes following the end of the session, mice received an IP injection of lithium chloride (180mg/kg). The following day (day 7) mice were placed back in the chamber for 30 minutes, with water freely available, and received an IP injection of saline vehicle 5 mins after the session ended. On day 8, mice underwent a second round of CTA and were placed in the chamber for 30 minutes with access to vanilla milk only (to encourage consumption of milk, which had already undergone CTA). Mice were given LiCl (180mg/kg) IP injection 5 minutes following the end of the session. Day 9 mirrored day 5, with vCA1 activity being imaged while mice received vanilla milk, low NaCl, and high NaCl solutions. Finally, on day 10, liquid preference was tested again as mice were given free access to vanilla milk and water for 30 mins.

Throughout training, animals were water restricted to ∼85-90% ad lib weight. A total of 10 distinct FOVs from 5 mice were included in the data set.

#### Experiment 6: Associative learning

Four neutral odors served as conditioned stimuli cues (heptaldehyde, - limonene, alpha-pinene, and ethyl butyrate, 2 sec) with cue contingencies counterbalanced across mice. Presentation of the CS+ cue was followed by a 2s trace period and subsequent delivery of US (reward US = ∼2 µl 10% sucrose solution; low shock US = 0.125mA amplitude, 250ms duration; high shock = 0.5mA amplitude, 250ms duration). No US was presented following the presentation of the CS-cue. Animals were not punished for off-target licking. Shocks were delivered via custom-made tail cuffs (one for each shock intensity) driven by precision animal shockers (Coulbourn Instruments, Holliston, MA). Trials were pseudorandomly interleaved, and each session consisted of 30 trials of each of the four trail types. The inter-trial interval between subsequent cues was chosen as a random sample from a uniform distribution between 17.5 and 27 seconds. Only the first 10 trials of each trial type were included for analysis.

Animals were trained on this paradigm for ∼7 consecutive days, with day 1 labeled “Early” and the final day of training “Late.” Throughout training, animals were water restricted to ∼85-90% ad lib weight. A total of 6 distinct FOVs from 6 mice were included in the data set.

### 1-photon imaging

3 weeks after surgery, mice were checked for GCaMP expression with a miniaturized microscope (Inscopix, Palo Alto, CA) and procedures previously described (Resendez et al., 2016). Mice were briefly anesthetized with 1.5% isoflurane at 1 L/min oxygen flow, and head-fixed into a stereotactic frame. The protective rubber mold was removed from the lens, and a magnetic baseplate was attached to a microscope and lowered over the implanted GRIN lens to assess the FOV for GCaMP+ neurons. If GCaMP+ neurons were visible, the baseplate was dental cemented in place onto the mouse headcap to allow for re-imaging of the same FOV for several weeks. Once baseplated, the same microscope was used for every imaging session with that mouse, and the focal plane on the hardware of the miniscope was not altered throughout the imaging experiments to ensure a constant FOV across sessions.

Ca2+ videos were recorded and processed with nVista acquisition and preprocessing software, version 1.1 (Inscopix, Palo Alto, CA), and triggered with a TTL pulse from EthoVision XT 10 and Noldus IO box system to allow for simultaneous acquisition of Ca2+ and behavioral videos for EPM and Arduino for the head-fixed behaviors. Ca2+ videos were acquired at 15 frames per second with 66.56 ms exposure. An optimal LED power was selected for each mouse based on GCaMP expression in the FOV (pixel values), and the same LED settings were used for each mouse throughout the series of imaging sessions, only a single FOV was imaged per animal. Videos were downsampled to 5 frames per second, cropped, and band-passed filtered as needed before Inscopix-based turboreg motion correction relative to a single reference frame.

### 2-photon imaging

Genetically encoded calcium imaging of GCaMP6f or 8m was used to assess the functional activity of individual neurons. Images were captured using an Ultima IV laser scanning microscope (Bruker Nano, Middleton, WI, acquired with Prairie View 5.4) equipped with resonant scanning mirrors and high-speed scan electronic controller, dual GaAsP PMTs (Hamamatsu model 7422PA-40), and motorized z focus (100 nm step size). GCaMP signal was filtered through an ET-GFP (FITC/CY2) filter set. Laser signal was provided by a MaiTai DeepSee mode-locked Ti:Sapphire laser source (Spectra-Physics, Irvine, CA) providing > 150kW max output at 920 nm. Acquisition speed was 30 Hz for 512 x 512 pixel images. Images were averaged 8x offline, yielding a final frame rate of 3.75 Hz.

Prior to each conditioning session, the imaging field of view (FOV) was determined, and imaging was conducted at that FOV for the entire session. For animals with multiple FOVs across sessions, each FOV was separated by > 100 µm in the z-dimension (dorsal-ventral) to ensure no overlap of cells across different FOVs. To facilitate re-identification of a specific FOV across sessions, the top of the GRIN lens served as a reference z-plane. Optimal laser power was determined for each FOV based on GCaMP expression level and was kept constant across sessions for a specific FOV. For each trial, imaging began 5 sec prior to stimulus onset and was terminated 7 sec after the US presentation.

In a subset of animals (n=8), two FOVs were collected using an objective z-piezo stage (Bruker, Middleton, WI) that rapidly (∼80ms) switched between different z-planes. To ensure each FOV consisted of distinct neuronal populations, each z-plane was separated by >100 µm and were manually inspected prior to recording. Recordings were averaged offline to a final frame rate of 3 Hz in order to closely match recordings that did not utilize piezo imaging.

### Signal extraction and cross-session registration

Videos were motion corrected offline using non-rigid motion correction based on template matching (Suite2p (Pachitariu et al., 2017)). Cell segmentation and calcium transient time series data were extracted using Constrained Non-negative Matrix Factorization for microEndoscopic data (CNMFe), a semi-automated algorithm optimized for GRIN lens Ca2+ imaging to denoise, deconvolve and demix calcium imaging data (Zhou et al., 2018). Putative neurons were manually inspected for appropriate spatial properties and Ca2+ dynamics and were visually checked against the corresponding motion corrected video in ImageJ. Ca2+ transient events were extracted using the OASIS algorithm (Friedrich et al., 2017) embedded within CNMFe. We used these inferred calcium events for all analyses, unless otherwise noted. Denoising (CNMFe) and deconvolution (OASIS) steps were applied.

Registration of cells across sessions imaged at the same FOV used probabilistic modeling of similarities between cell pairs across sessions (CellReg, (Sheintuch et al., 2017)). Briefly, spatial footprint maps were generated for each session by projecting the spatial filter of each cell onto a single image. Spatial footprint images from sessions imaged at the same FOV were then aligned. The distribution of similarities between pairs of neighboring cells were subsequently modeled via centroid distance to obtain an estimation for their probability of being the same cell (P_same_). Cells were then registered across sessions via a clustering procedure that utilizes the previously obtained probabilities, with a probability threshold of 0.8. The average P_same_ value for registered cells was 0.95. All putative matches were visually inspected.

### Data analysis

For statistical analyses and figures (aside from activity heatmaps), calcium event activity was separated into 2-second bins and average activity during each bin was used. All statistical analyses were two-sided. Data distribution was assumed to be normal but this was not formally tested. No statistical methods were used to pre-determine sample sizes but our sample sizes are similar to those reported in previous publications (Biane et al., 2023). For all figures: * p< 0.05, ** p < 0.01, *** p < 0.001. See Supplementary Table 1 for all statistical analysis details.

#### Population decoding

A linear decoder was used to discriminate activity patterns into two discrete categories (Christopher Bishop, n.d.):

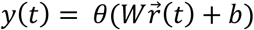

where *y* is the predicted label of the population activity pattern 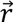 recorded at time 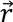 and takes two values corresponding to two classes of patterns to decode (for example, two odor identities), *W* is the vector of weights assigned to each cell, and *b* is a bias term constant. Decoding parameters were attained via a supervised learning protocol with labeled data and used a support-vector machine (SVM) with a linear kernel (python/scikit/linearSVC) and regularization (C) set to 1 (ie, default value). Results are reported as the generalized performance of the decoder using cross-validation. When multiple categories were involved, e.g., more than two trial types, multiple linear decoders were trained on pairs of discrete categories combined using majority-based error-correction codes. Testing of different regularization and kernel parameters was executed with python/scikit/svm/SVC, using default settings of any additional parameters.

We defined the patterns of calcium activity by computing the mean event rates for each individual cell during two-second time bins. Pseudo-population recordings were generated by combining cell data across multiple animals/FOVs. For decoding, one-half of trials were randomly selected from each class and pseudo-population activity from these trials was used to train the decoder, while the remaining held-out half was used to evaluate the decoder’s generalization performance. When comparing decoding accuracy between neural populations of different size, we trained our decoder on a random subsample of cells from the more numerous population equal to that of the smaller population. We repeated the operation 100 times and then combined the cross-validated decoding accuracies of all random choices together to get a single sample of decoding accuracies (i.e., single data point reflecting the mean of all 100 iterations). We repeated the procedure 10 times to perform statistical comparisons across groups and against chance performance. A two-sided Mann-Whitney U Test was used to compare decoding accuracies between groups, and Bonferroni correction used for multiple comparisons.

For decoding against baseline, we used population activity during the 2-second time bin that began four seconds prior to stimulus onset as baseline data. Cross-session decoding followed the same procedure as within-session decoding. In the case of cross-session decoding, only cells registered across the compared sessions were included.

#### Multidimensional Scaling (MDS)

We performed 2-dimensional MDS scaling of event data using python/scikit/MDS. As with decoding, we used all cells recorded across all mice to produce one pseudopopulation. The Euclidean distance was taken between each trial type, and this process was repeated 10 times. Bar charts of Euclidean distance show the mean ± SEM of all runs.

#### Mahalanobis distance

To examine the Mahalanobis distance between population activity patterns occurring during presentations of different stimuli, we first randomly downsampled the number of neurons to match the number of trials (this was necessary to minimize failures when attempting to invert the covariance matrix). We next estimated the covariance matrix using python/numpy/cov. The inversion of this matrix was calculated using python/numpy/linear algebra/inv. If the inversion failed, Ledoit-Wolf regularization was used (python/scikit learn/covariance/LedoitWolf). Finally, the Mahalanobis distance (M) was computed using python/scipy/spatial/distance/Mahalanobis.

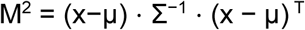

where x and μ are the mean population responses for each of the stimuli being compared, T is their transpose, and Σ is the covariance matrix. This process was repeated 50 times, and the mean of all 50 runs was returned. We repeated the above procedure 10 times, providing 10 data. Finally, data were z-scored with respect to a 2 sec time window prior to stimulus delivery.

#### Single-cell responsivity

Data used for heatmaps of calcium-traces or inferred events were not binned. For each cell, z-scores were computed over the entire dataset for a specific condition (e.g., sucrose reward trials). To identify cells whose activity was modulated during specific epochs (e.g., stimulus presentation), Ca2+ events were shuffled in time for each cell (1000 iterations), and stimulus rates were re-calculated from those time periods to generate a null distribution of stimulus Ca2+ event rates for each cell. A cell was considered stimulus responsive if its Ca2+ event rate during those periods exceeded a 1 SD threshold from the null distribution. The False Discovery Rate (FDR) was applied to correct for multiple comparisons. Cells with an adjusted p-value < 0.05 were classified as responsive.

#### Across-stimulus and across-session overlaps in single-cell responsivity

To compute overlapping responses, we determined the number of cells that showed statistically significant responses to a specific stimulus (e.g., peanut oil odor) or task epoch (e.g., cue period). To assess statistical significance, we pooled cells from all mice to generate pseudo-simultaneous recordings. Chance distributions were generated by randomly assigning responses to all cells for each of the two sessions being compared, with probabilities that matched the proportion of responsive cells for each session as in the real data. We obtained chance distributions by computing the overlap for each random assignment and repeating the procedure 10000 times. Statistical significance was assessed for the actual overlap between the two sessions by computing the probability of obtaining that value from the chance distribution assuming a normal distribution of estimated mean and variance.

#### Effect size estimation

For Mann Whitney U tests and Wilcoxon tests versus chance, effect size was determined using:

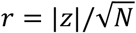

where *N* is the total sample size and

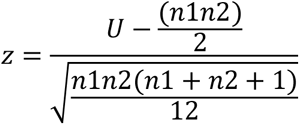

Where *n*1 is the sample size of sample 1, *n*2 the sample size of sample2, and *U* is the U test statistic obtained from the statistical test output. For t-test analysis, Cohen’s d was used, defined as the difference between group means, divided by their pooled variance. For Fishers analysis, the odds ratio was obtained directly from the test output. For one-way ANOVA, ETA^2 was obtained directly from the test output.

### Data Availability

All source data can be downloaded from the Kheirbek lab github site (github.com/mkheirbek)

### Code Availability

The analysis code supporting this study is available from the Kheirbek lab github site (github.com/mkheirbek).

### Contact for reagents and resource sharing

Further information and requests for resources and reagents should be directed to M.A.K.

